# Peptidoglycan editing provides immunity to *Acinetobacter baumannii* during bacterial warfare

**DOI:** 10.1101/2020.04.23.058008

**Authors:** Nguyen-Hung Le, Katharina Peters, Akbar Espaillat, Jessica R. Sheldon, Joe Gray, Gisela Di Venanzio, Juvenal Lopez, Bardya Djahanschiri, Elizabeth A. Mueller, Seth W. Hennon, Petra Anne Levin, Ingo Ebersberger, Eric P. Skaar, Felipe Cava, Waldemar Vollmer, Mario F. Feldman

## Abstract

Peptidoglycan (PG) is essential in most bacteria. Thus, it is often targeted by various assaults, including the host immune response, antibiotic treatment and interbacterial attacks via the type VI secretion system (T6SS). Here, we report that the Gram-negative bacterium *Acinetobacter baumannii* strain ATCC 17978 produces, secretes and incorporates the non-canonical D-amino acid D-Lysine into its PG during stationary phase. We show that PG editing increases the competitiveness of *A. baumannii* during bacterial warfare by providing immunity against peptidoglycan-targeting T6SS effectors from various bacterial competitors. We propose that PG editing has evolved as an effective strategy for bacteria to overcome T6SS attacks. In contrast, we found that D-Lys production is detrimental to pathogenesis due, at least in part, to the activity of the human enzyme D-amino acid oxidase (DAO), which degrades D-Lys producing H_2_O_2_ toxic to bacteria. Phylogenetic analyses indicate that the last common ancestor of *A. baumannii* possessed the ability to produce D-Lys. However, this trait was independently lost multiple times, likely reflecting the evolution of *A. baumannii* as a human pathogen.

**One sentence summary:** *Acinetobacter baumannii* attains immunity against nonkin competitors during T6SS warfare by incorporating D-Lysine into its peptidoglycan.

## Main Text

Peptidoglycan (PG) is a major component of the bacterial cell envelope. Layers of PG surround the cytoplasmic membrane, maintaining cell shape and providing resistance to osmotic stress. PG is composed of glycan chains of alternating *N*-acetylglucosamine (Glc*N*Ac) and *N*-acetylmuramic acid (Mur*N*Ac) that are connected through Mur*N*Ac-attached peptides (*1*). Despite its rigidity, PG is necessarily a dynamic structure, requiring constant turnover to enable fundamental processes such as growth and cell division (*2*). PG synthesis is a multi-step process. The PG precursor lipid II, which consists of a Glc*N*Ac-Mur*N*Ac pentapeptide (L-Ala-D-iGlu-mDAP-D-Ala-D-Ala in Gram-negative bacteria) linked to a lipid carrier, is synthesized at the cytoplasmic membrane and subsequently translocated into the outer leaflet of the cytoplasmic membrane. Glycan chains are then polymerized by glycosyltransferases and peptide cross-links are formed through transpeptidation reactions catalyzed by DD-transpeptidases and LD-transpeptidases.

Due to its essentiality, the bacterial cell wall is targeted by various potentially fatal threats, including attacks from bacterial competitors. Competition among Gram-negative bacteria is largely mediated by the type VI secretion system (T6SS) (*3*). The T6SS is a dynamic nanomachine structurally related to contractile phage tails, and it is employed to deliver toxic effector proteins, including PG hydrolases, from an attacking cell (predator) to nearby competitors (prey) in a contact-dependent manner (*4–6*). Immunity to T6SS-dependent killing is accomplished by the expression of immunity proteins, which specifically bind and inactivate their cognate effector. The limited repertoire of immunity proteins encoded by one bacterium is insufficient to protect against the large diversity of T6SS effectors encoded by bacterial predators. Therefore, the primary role of immunity proteins is to avoid lethal interactions between sister cells (*7–9*). General resistance mechanisms to prevent T6SS-dependent killing by nonkin competitors remain poorly characterized (*10*).

Additional threats to the integrity of the bacterial cell wall include the host immune system, antibiotics, and osmotic stress (*11–13*). Constant exposure to these threats has exerted selective pressure favoring bacterial species capable of modifying their PG to evade or withstand fatal threats. These modifications can occur during PG precursor synthesis or after PG precursors are incorporated into the growing cell wall meshwork (*14*). PG modifications enabling immune evasion and resistance to antibiotics have been extensively studied and generally include chemical modifications to the glycan backbone or changes in pentapeptide composition or degree of crosslinking (*12*, *14–18*). Additionally, PG editing with non-canonical D-amino acids (NCDAAs) has been shown to make bacteria more resistant to osmotic stress (*19*). NCDAAs have been best characterized in *Vibrio cholerae*, which modifies up to ~5% of its PG subunits with D-Met during stationary phase. In addition to incorporating D-Met into its PG, this organism produces and secretes a wide variety of NCDAAs to millimolar concentrations (*19*). Importantly, some of the secreted NCDAAs inhibit the growth of diverse bacteria at physiological concentrations, making NCDAAs important mediators of interbacterial competition (*20*).

The Gram-negative bacterial pathogen *Acinetobacter baumannii* has a remarkable ability to withstand antibiotic treatment and persist in healthcare settings. As such, it is categorized by the World Health Organization (WHO) and the U.S. Centers for Disease Control and Prevention (CDC) as a critical priority for the research and development of novel antimicrobial therapies. Given the remarkable ability of *A. baumannii* to withstand a wide variety of environmental stressors (*21*), we investigated whether this organism modifies its PG during stationary phase, as has been reported in other bacterial species (*1*, *19*, *22*). To this end, we purified PG from Ab17978 grown to exponential (PG_exp_) or stationary phase (PG_stat_), digested it with muramidases, and analyzed the resulting muropeptides by HPLC. The muropeptide profile obtained from PG_exp_ is similar to that of *E. coli* (Figures 1A, S1 and S2). However, the HPLC chromatogram derived from PG_stat_ showed five unique peaks (Figure 1A). Mass spectrometry analysis identified these products as monomeric or cross-linked muropeptides containing a lysine residue (Lys) in the fourth (L-Ala-D-iGlu-mDAP-Lysine, hereafter “Tri-Lys”) or fifth (L-Ala-D-iGlu-mDAP-D-Ala-Lysine, hereafter “Tetra-Lys”) position (Figure 1B and S8, Table S1). Quantification of HPLC profiles showed that the Tri-Lys muropeptide and its cross-linked forms constitute ~24% of the total muropeptides of PG_stat_ (Figure 1A, Table S2). Lys-containing muropeptides were also present in PG_exp_ but in a considerably lower amount (~6% of total muropeptides). In contrast, Tetra-Lys muropeptides were unique to PG_stat_ and accounted for ~10% of the total muropeptides. Together, our results indicate that in stationary phase, Ab17978 is capable of modifying about one-third of its PG with Lys.

**Fig. 1.**
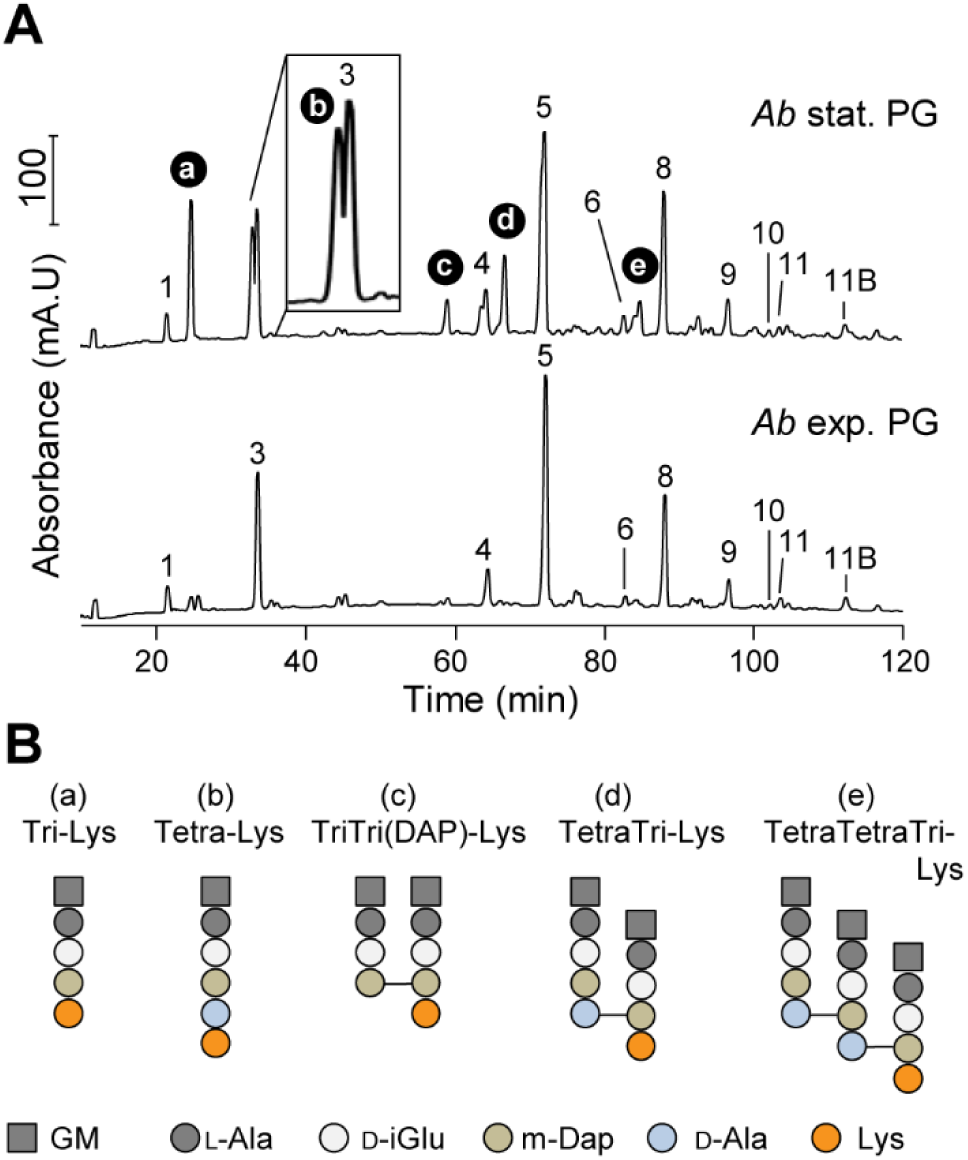
Ab17978 incorporates D-Lys into its PG at stationary phase. **(A)** Chromatograms of purified muropeptides from Ab17978 during stationary (stat.) and exponential (exp.) phase. Main muropeptides (No. 1-11B) are numbered according to Boll *et al*., 2016. D-Lys-containing muropeptides (a-e) were identified by mass spectrometry. Proposed muropeptide structures are shown in Fig. S2. **(B)** Proposed structures of D-Lys-modified muropeptides a-e. GM, *N*-acetylglucosamine-*N*-acetylmuramitol; L-Ala, L-alanine; D-iGlu, D-isoglutamate; m-Dap, meso-diaminopimelic acid; Lys, lysine; D-Ala, D-alanine. “Tri” or “Tetra” indicate the number of amino acids in the peptides chain preceding Lys.

Mass spectrometry is unable to differentiate L-Lys from D-Lys. Previous work showed that bacteria that produce and incorporate NCDAAs into their PG meshwork also secrete NCDAAs into the culture supernatant at millimolar concentrations (*19*, *22*). Thus, D-Lys secretion by Ab17978 would suggest editing with the D-enantiomer. We developed a colorimetric assay using the D-Lys-specific oxidase AmaD from *Pseudomonas putida* to detect and quantify the concentration of D-Lys in culture supernatants of Ab17978 (Figure 2A) (*23*). Using our AmaD assay, we determined that Ab17978 produces and secretes D-Lys into culture supernatants to a concentration of ~0.3 mM (Figure 2B), which is comparable to levels of NCDAA secreted by other bacterial species (*19*, *24*). D-Lys secretion has not been reported in *Acinetobacter* leading us to investigate how D-Lys is produced. Amino acid racemases are enzymes that catalyze the conversion between L-amino acids and D-amino acids. Ab17978 encodes five putative amino acid racemases, only one of which is predicted to have a signal peptide (ACX60_11360, hereafter “RacK”). To identify the racemase involved in D-Lys production, we constructed mutant strains of Ab17978 lacking each racemase and determined the levels of D-Lys in their culture supernatants. Deletion of only one racemase, RacK, completely abrogated D-Lys secretion (Figure S3). Importantly, complementation of the *racK* gene (RacK+) restored D-Lys secretion to levels comparable with wild-type (WT) Ab17978 (Figure 2B). Having identified RacK as responsible for D-Lys production, we determined if D-Lys was the Lys enantiomer incorporated into the PG of Ab17978 during stationary phase. We analyzed muropeptides from Ab17978 WT, RacK and RacK+ by HPLC. Indeed, PG_stat_ from RacK lacked the peaks corresponding to Tri-Lys and Tetra-Lys subunits, all of which were present in the WT and RacK+ strains (Figure 2C). Together, our results demonstrate that Ab17978 secretes D-Lys to millimolar concentrations and that PG editing is dependent on the production of D-Lys by RacK. Because RacK is predicted to localize to the periplasm, we propose that PG editing with NCDAAs in Ab17978 occurs in the periplasm, as has been suggested for *V. cholerae* (*19*, *22*). Previous transcriptomics study in Ab17978 showed that *racK* was also upregulated in biofilms compared to cells in exponential phase (*25*).

**Fig. 2.**
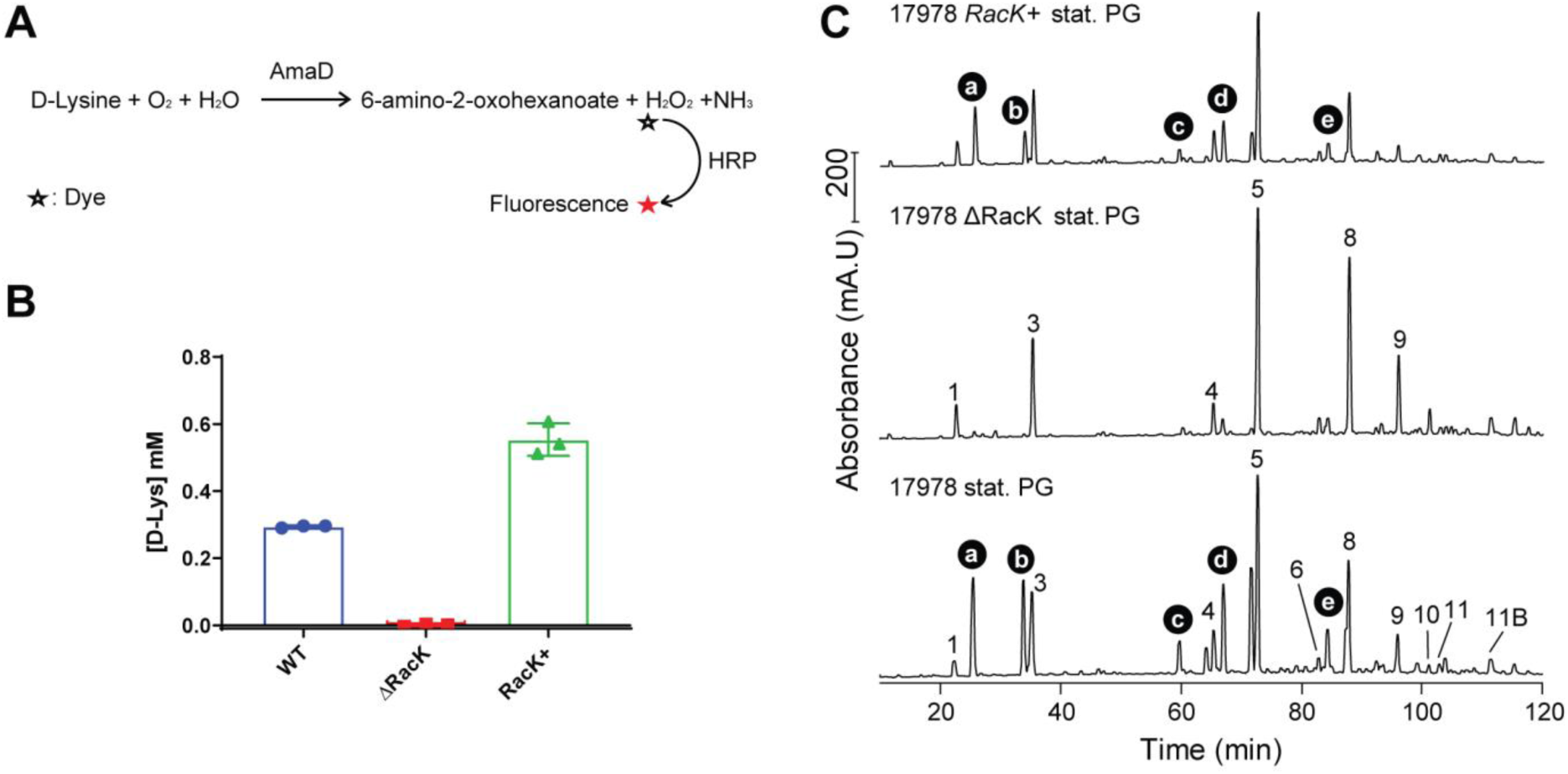
Racemase RacK is responsible for D-Lys secretion and PG editing. **(A)** Overview of the colorimetric assay used to quantify D-Lys secretion. Purified D-Lys oxidase AmaD from *Pseudomonas putida* catalyzes the oxidative deamination of D-Lys to release 6-amino-2-oxohexanoate, hydrogen peroxide (H_2_O_2_) and ammonium (NH_3_). H_2_O_2_ produced was then quantified by treatment with horse radish peroxidase (HRP) and a fluorophore dye. The amount of fluorescence is converted to D-Lys concentration through a standard curve **(B)** Quantification of D-Lys present in supernatant fractions of *Ab*17978 WT, 17978ΔRacK and the complemented strain (17978 RacK+). Data represent mean ± SD of 3 biological replicates. **(C)** Chromatograms of purified, stationary phase muropeptides from the indicated strains. Muropeptide structures are shown in Fig. S2

Virtually all bacteria have the ability to incorporate NCDAAs, either produced by their own racemases or present in the environment, into their PG meshwork (*14*, *26*, *27*). This can be accomplished by the activity of DD- or LD-transpeptidases (*22*, *27–29*). Previous work has demonstrated that PG editing with NCDAAs impacts cell physiology and decreases both DD- and LD-crosslinks, likely by competing with transpeptidase activity (*19*, *22*, *28–30*). Consistently, PG of Ab17978 WT and RacK+ showed a lower percentage of muropeptides in crosslinking compared with Ab17978 ΔRacK, indicating that D-Lys interferes with the PG cross-linking activity of transpeptidases (Table S4). Previous studies have shown that PG crosslinking enhances a bacterium’s ability to withstand stress (*31*, *32*). Thus, we next tested whether PG editing with D-Lys affected the ability of Ab17978 to withstand a variety of environmental stressors. We found that Ab17978 RacK was nearly indistinguishable from WT in growth under pH and osmotic stress as well as cell morphology (Figure S4), which suggest an alternative role for D-Lys in *Acinetobacter*. NCDAAs are considered part of the warfare between bacterial species, as certain secreted D-amino acids can inhibit the growth of diverse bacteria (*20*). However, we found that the concentration of D-Lys present in the spent media of Ab17978 was not sufficient to inhibit the growth of *P. aeruginosa* PA01, *A. nosocomialis* strain M2 (M2) or *A. baumannii* strain 19606 (Ab19606) (Figure S5).

We then hypothesized that PG editing with D-Lys could constitute a protective mechanism against T6SS-mediated attacks from nonkin bacteria. In this model, PG editing masks the target of T6SS PG hydrolase effectors, similarly to how bacteria become resistant to antibiotics by modifying the structure of the antibiotic target. The H1-T6SS of *P. aeruginosa* PA01 delivers at least two PG hydrolase effectors into target cells: the amidase (or D,L-endopeptidase) Tse1 and the muramidase Tse3 (*4*). Thus, to test our hypothesis we performed bacterial competition assays with *P. aeruginosa* PA01 as the predator and Ab17978 WT or ∆RacK as prey. We found that ∆RacK was ~15-fold more susceptible to T6SS killing compared with WT Ab17978 (Figure 3A). Consistently, UPAB1 RacK+ showed ~25-fold higher survival than WT UPAB1 when co-incubated with PA01 (Figure 3A and Figure S6). Similarly, we observed that PG editing by D-Lys increased the survival of *A. baumannii* against *Serratia marcescens* and *A. nocosomialis* M2 (Figure 3A). Collectively, these results indicate that PG editing with D-Lys provides *A. baumannii* immunity against T6SS-dependent attacks from diverse bacteria.

**Fig. 3.**
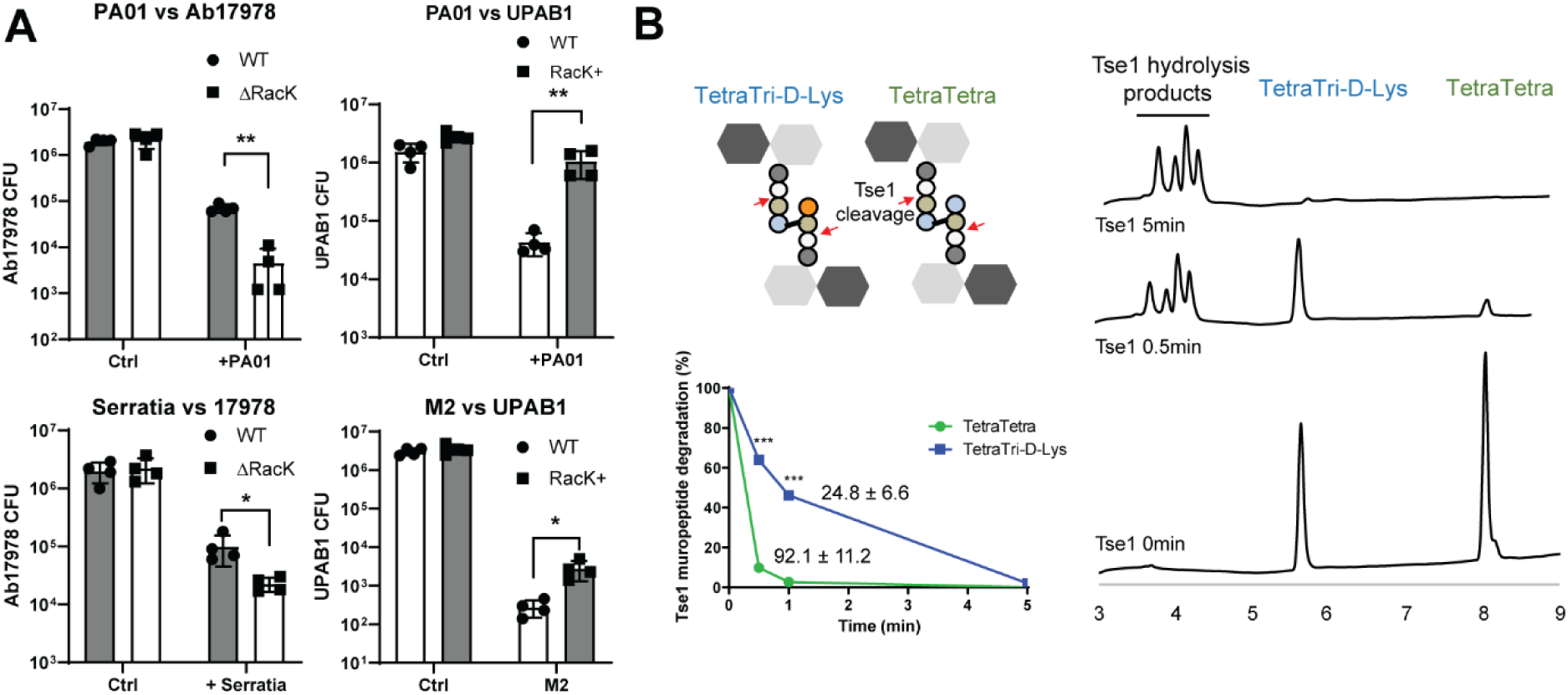
(A) PG editing with D-Lys provides protection to T6SS-dependent attack from *P. aeruginosa* PA01. Competition assay between PA01, *S. marcescens* (Serratia) or A. nocosomialis M2 as predator and *A. baumannii* strains expressing (Ab17978 WT or UPAB1 RacK+) or not expressing RacK (Ab17978ΔRacK or UPAB1 WT) as prey. Bar graphs represent the mean ± SD from 4 biological replicates. **(B)** Non-edited (TetraTetra) and edited muropeptides (TetraTri-D-Lys) were treated together with purified PA01 amidase effector Tse1. Reaction products were analyzed by HPLC at time t = 0 (Ctrl), t = 0.5 min and t = 5 min. Undigested muropeptides were quantified and specific activity of Tse1 against TetraTetra and TetraTri-D-Lys was calculated. Specific activity values shown on graph are expressed in mg of muropeptide degraded/min/Tse1 concentration. Data present mean ± SD of 2 replicates. Statistical analyses were performed using the Student’s unpaired t test, *p < 0.05, **p<0.01.

To gain better insight into the mechanism of PG editing-based immunity to T6SS-mediated attacks, we compared the activity of *P. aeruginosa* PA01 Tse1 towards non-edited (Tetra-Tetra) and D-Lys-edited (TetraTri-D-Lys) dimeric muropeptides. A mix of both muropeptides was incubated with purified Tse1 and samples were taken at different time points and analyzed by HPLC (Figure 3B). At 0.5 min, we found that most of the TetraTetra muropeptides were hydrolyzed by Tse1, whereas nearly 40% of TetraTri-D-Lys remained uncleaved. Importantly, we observed a ~4-fold decrease in the specific activity of Tse1 toward the D-Lys-modified muropeptide compared with the non-edited muropeptide, consistent with the results of our bacterial competition assays (Figure 3A). Together, these results support the hypothesis that PG editing in Ab17978 provides an immunity protein-independent mechanism to prevent T6SS-dependent killing by masking the target PG from the injected effector.

Some bacterial species modify their PG to counteract the host innate immune response (*12*, *33*). We examined whether the presence of RacK would affect *A. baumannii* pathogenesis in the murine acute pneumonia model of infection (*34*). Ab17978 or UPAB1 strains either encoding or lacking RacK were introduced by intranasal inoculation, and bacterial burdens were enumerated 36 h post infection. Unexpectedly, strains unable to produce D-Lys showed significantly higher bacterial burdens (1 to 2 log CFU) in the kidney, heart and liver, suggesting that D-Lys production is disadvantageous for bacterial dissemination (Figure 4A). Recent findings have suggested that the D-amino acid oxidase (DAO), expressed by human neutrophils, macrophages and epithelial cells, contributes to host defense against various pathogens through the production of H2O2, a by-product of the DAO-dependent oxidative deamination of D-amino acids (*24*, *35*, *36*). Thus, we hypothesized that D-Lys-producing bacteria would be more susceptible to DAO-dependent killing. To test this, we incubated stationary phase Ab17978 or UPAB1 strains either encoding or lacking RacK with purified DAO and enumerated surviving CFU after 4.5 h. DAO showed significant antibacterial activity against *A. baumannii* strains harboring RacK, resulting in a 103-fold reduction in CFU compared with strains not producing D-Lys (Figure 4B). Importantly, addition of purified catalase, an enzyme that eliminates H_2_O_2_, provided full protection against DAO, indicating that *A. baumannii* killing by DAO is H_2_O_2_-dependent. Together, our results demonstrate that D-Lys production is detrimental to pathogenesis, at least in part due to DAO activity.

**Fig. 4.**
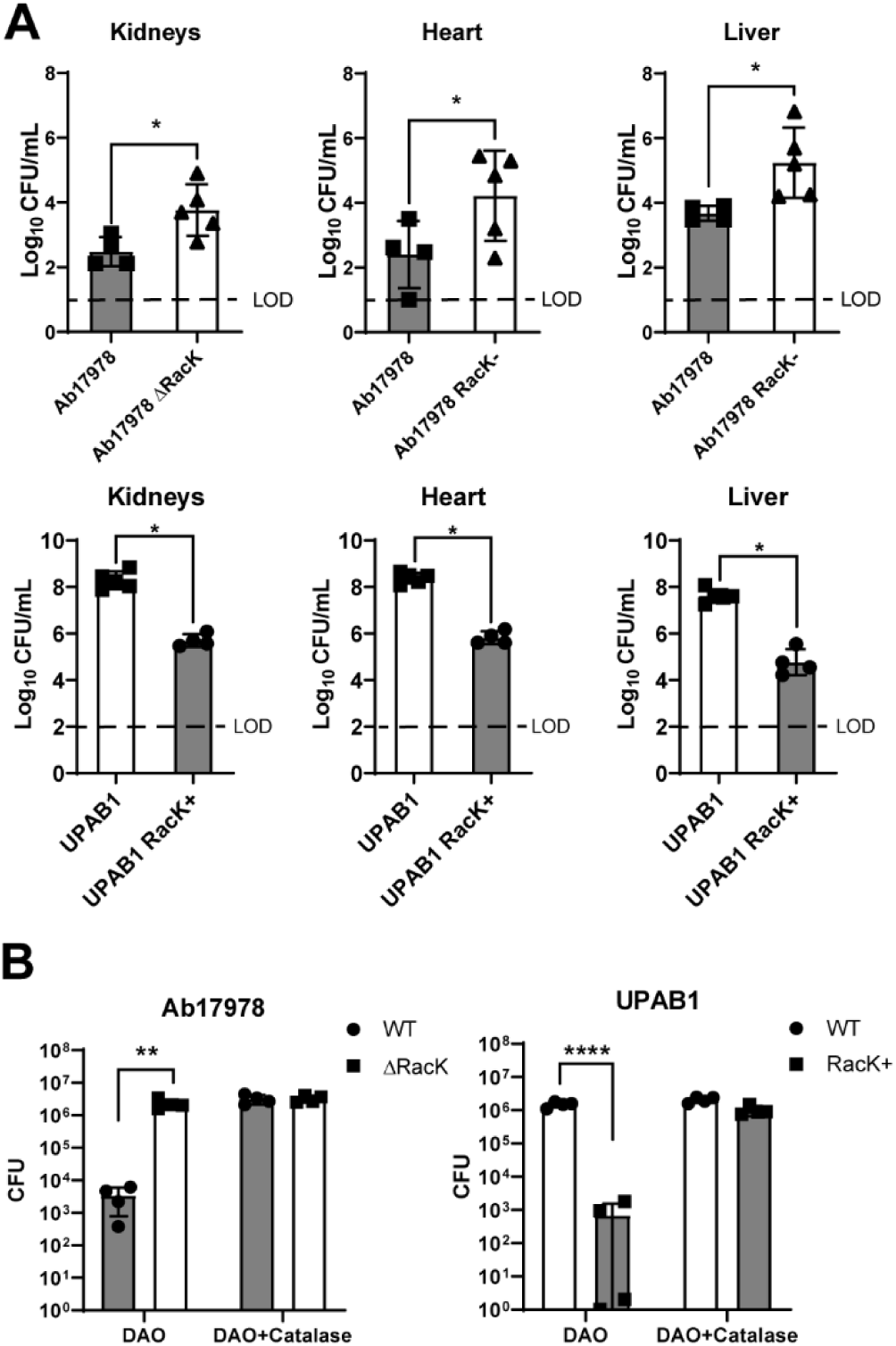
(A) RacK is detrimental in a murine model of acute pneumonia. Mice were intranasally inoculated with either the Ab17978 WT or ΔRacK (left panel), or the UPAB1 WT or RacK+ (right panel). Following 36 h of infection, the bacterial burdens of the kidneys, heart, liver, spleen, lungs and blood were determined. Each symbol represents an individual mouse. Statistical analyses were performed using the Mann–Whitney U test, and LOD represents the limit of detection. **(B) DAO has increased antibacterial activity on *A. baumannii* strains having RacK.** WT and ΔRacK Ab17978 (left panel) or WT and RacK+ UPAB1 (right panel) were incubated with DAO or DAO+Catalase for 4.5h and CFU were enumerated. Bar graphs represent the mean ± SD of 4 biological replicates. Statistical analyses were performed using the unpaired Student’s t test, *p < 0.05, **p< 0.01, ****p< 0.0001.

The benefit of PG editing in bacterial competition and detriment to pathogenesis present an interesting evolutionary paradox, making it difficult to predict whether natural selection would favor *Acinetobacter* strains capable or incapable of modifying their PG. To this end, we performed a phylogenetic profiling analysis of RacK across 3,052 *Acinetobacter* strains representing 53 species. We found that RacK is largely confined to the *Acinetobacter calcoaceticus/baumannii* (Acb-) complex, occurring only sporadically in other species (Figure S7). Within the Acb-complex, RacK is prevalent in the two earliest branching lineages, suggesting its presence in the last common ancestor of the Acb-complex (Figure 5). Furthermore, RacK is always flanked by the same four genes, and this gene order is maintained irrespective of the presence of RacK (Figure 5). A phylogenetic analysis provided no evidence that the evolutionary history differs between the four flanking genes and RacK, indicating that the five genes were likely present in the last common ancestor of *A. baumannii*. Importantly, we found that only 7% of *A. baumannii* strains encode RacK and these strains are distributed across different clades in the *A. baumannii* phylogeny (Figures 5 and S6). Thus, our results indicate that RacK was encoded by the last common ancestor of the Acb-complex and was lost multiple times independently during *A. baumannii* evolution.

**Fig. 5.**
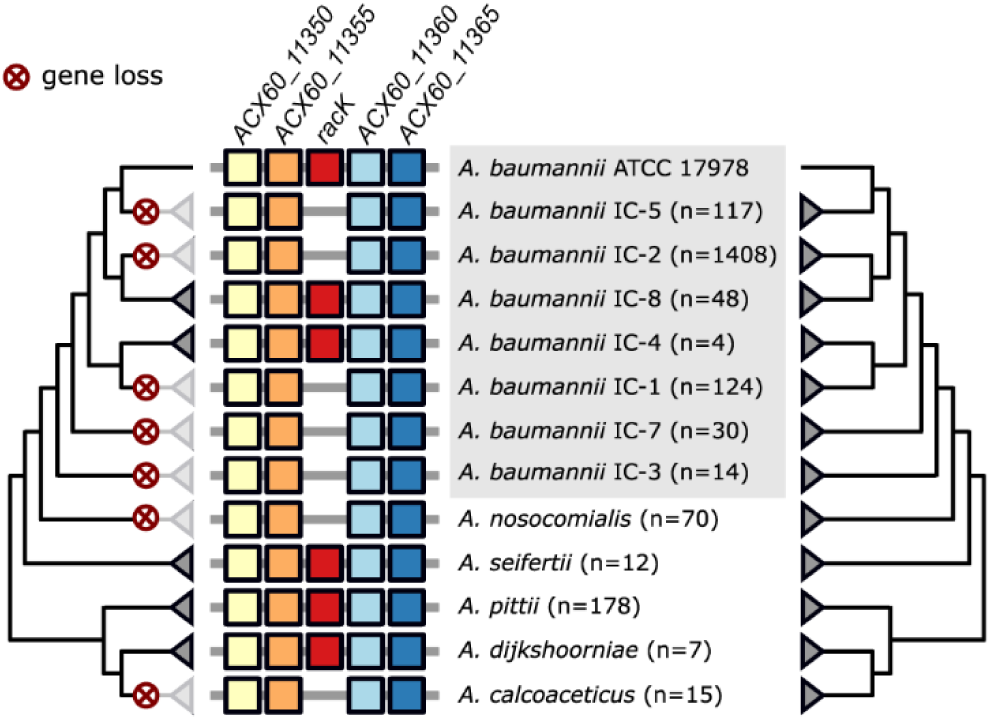
The evolutionary history of A. baumannii RacK. The presence of RacK (red boxes) is limited to individual lineages within A. baumannii, while it is ubiquitously present in the closely related species A. seifertii, A. pitti, and A. dijkshoorniae. Whenever present, RacK is embedded within the same four flanking genes (coloured boxes) throughout the Acb complex. This gene order is maintained in the species and strains were RacK is missing. The phylogenetic relationships of RacK (left tree) and of the four flanking genes (right tree) is congruent, indicating that the contemporary patchy distribution of RacK is a consequence of recurrent and lineage specific gene loss (marked by the red crosses).

Our results are summarized in Fig 6. During stationary phase, the periplasmic racemase RacK catalyzes the conversion of L-Lys to D-Lys. Once produced, D-Lys is incorporated into the PG of Ab17978 by Ldt and penicillin binding proteins (PBPs) (*22*, *28–30*). PG editing with D-Lys confers immunity to T6SS-dependent attacks by reducing the specific activity of PG hydrolase effectors, such as Tse1. Because T6SS effectors recognize and cleave conserved amide bonds in PG, we propose that certain types of PG editing constitute an immunity mechanism against T6SS-dependent killing. Indeed, previous work reported that L-DAP amidation in the PG of the environmental bacterium *Gluconobacter frateurii* resulted in reduced *in vitro* digestion by T6SS endopeptidases compared with non-modified PG (*18*). This mechanism of T6SS immunity is independent of immunity proteins, which bind with high specificity to their cognate T6SS effectors. In contrast, PG editing-mediated immunity might protect against various T6SS effectors that recognize canonical PG structures. We therefore propose that PG editing provides *A. baumannii* with a type of “innate immunity” against diverse bacterial competitors. It is tempting to speculate that additional common targets for T6SS effectors (or T7SS effectors in Gram-positive bacteria) are also modified to evade the toxicity of these effectors. However, PG modification may not protect against all bacteria, as the degree of protection may depend on the specific effectors secreted by different species.

**Fig. 6.**
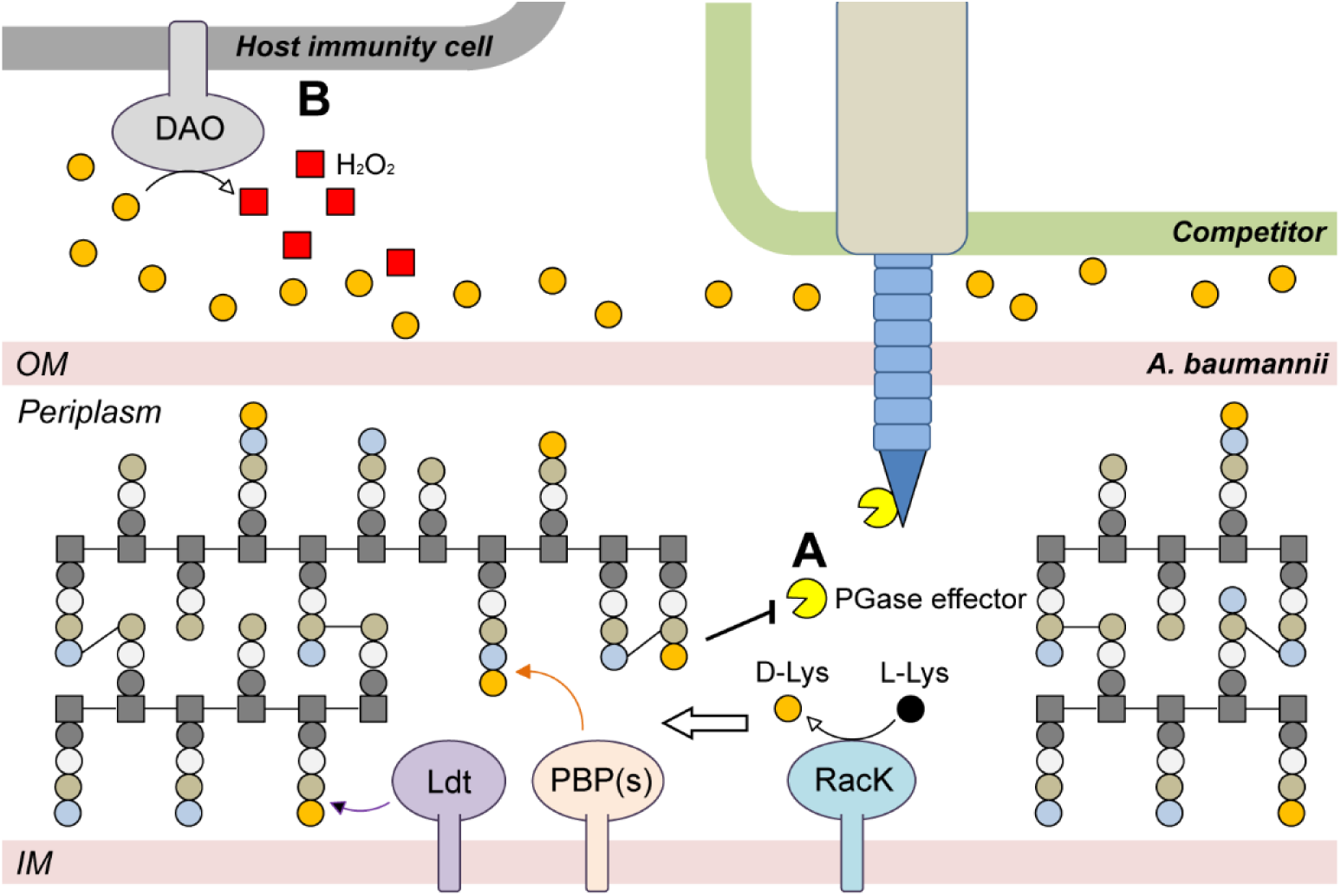
Proposed model of D-Lys production role in *Acinetobacter spp*. **(A)** The Lysine racemase RacK converts L-Lys to D-Lys in the periplasm. Presence of D-Lys in the periplasmic space leads to PG editing by Ldts and PBPs in competition with their canonical cross-link activity. This PG editing mechanism provides a form of innate immunity against foreign PG targeting T6SS. **(B)** As a trade-off, secretion of D-Lys during infection increase host DAO activity and thus decrease *A. baumannii* virulence.

Our data show that D-Lys production is unfavorable for *A. baumannii* pathogenesis. We have shown that, *in vitro*, purified DAO is able to recognize the D-Lys secreted by *A. baumannii* and produce enough H2O2 to kill bacteria expressing RacK. DAO is known to be produced by key players in the initial immune response against *A. baumannii* infections (*37–40*). Despite the competitive advantages conferred by the expression of RacK during interbacterial competition, modern clinical Ab strains as well as other *Acinetobacter* species of the Acb-complex have relinquished their copy of the *racK* gene. Indeed, strains from the most common lineages of clinical *A. baumannii* strains, such as international clone 1 (IC1), IC2, and IC3, do not carry the *racK* gene. This is reminiscent of the loss of the T6SS via genetic disruptions to the T6SS gene locus or the accumulation of inactivating point mutations in ~ 40% of *A. baumannii* clinical isolates (*41*, *42*). Both PG editing and a functional T6SS are crucial in the environment but appear to impose a fitness cost during infection. These examples illustrate the multiple adaptations that *A. baumannii* has undertaken to become a pathogen capable of successfully infecting the human host.

## Acknowledgements

This work was supported by National Institute of Allergy and Infectious Diseases (NIAID) Grant R21 AI137188. Work in the Vollmer lab was supported by the Wellcome Trust grant 101824/Z/13/Z and the United Kingdom Medical Research Council within the Antimicrobial Resistance Cross-Council Initiative collaborative grant MR/N002679/1. Work in the Cava lab was supported by the Swedish Research Council (VR), the Knut and Alice Wallenberg Foundation (KAW), the Laboratory of Molecular Infection Medicine Sweden (MIMS), and the Kempe Foundation. Work in the Skaar lab was supported by National Institute of Allergy and Infectious Diseases (NIAID) Grant R01 AI101171.

## Author Contributions

Conceived and designed the experiments: N.H.L., K.P., A.E., J.R.S., G.D.V., B.D., E.A.M., S.W.H., J.G., I.E., E.P.S., F.C., W.V. and M.F.F. Performed the experiments: N.H.L., K.P., A.E., J.R.S., G.D.V., J.L., B.D., E.A.M., J.G. and S.W.H. Analyzed the data: N.H.L., K.P., A.E., J.R.S., G.D.V., B.D., E.A.M., J.G., P.A.L., I.E., E.P.S., F.C., W.V. and M.F.F. Wrote the paper: N.H.L., J.L., and M.F.F.

## Data and materials availability

All data is available in the main text or the supplementary materials.

