## Supplemental Information for "Peptidoglycan editing provides immunity to *Acinetobacter baumannii* during bacterial warfare"

**This PDF file includes:**

Materials and Methods

Figs. S1 to S7

Tables S1 to S6

### Materials and Methods

#### *Bacterial strains and growth conditions*

Bacterial strains used in this study are listed in Supplementary Table S5. Unless otherwise noted, strains were grown in lysogeny broth (LB) liquid medium at 37 °C with shaking (200 rpm). The antibiotics rifampicin (150 µg per ml), irgasan (25 µg per ml), kanamycin (7.5 or 50 µg per ml), gentamycin (20 µg per ml), chloramphenicol (15 µg per ml), carbenicillin (100 or 200 µg per ml) and zeocin (50 µg per ml) were added when necessary. Spontaneous rifampicin-resistant mutant strains were obtained by plating an overnight culture on LB agar with rifampicin.

#### *PG isolation and analysis*

PG from exponential and stationary cells from *A. baumannii* strains was isolated and analyzed by HPLC as previously described (*42*, *43*). Briefly, *A. baumannii* strains were grown at 37°C in 500 mL of LB media to a final $OD_{600}$ of 0.6 for exponential and grown for 24h for stationary phase samples. Cells were collected by centrifugation for 15 min at 4°C and 7,000 × *g*. Cell pellets were resuspended in 50 mL cold water and lysed by drop wise addition to 50 mL boiling 8% SDS under vigorous stirring. Samples were boiled for further 30 min to ensure complete solubilization of the membranes and degradation of the high molecular weight DNA. Crude PG samples were collected by ultracentrifugation 60 min at 110,000 × *g* at 25°C. Pellets were washed several times with phosphate buffer (PB) pH 6.0 to remove SDS. Crude PG samples were treated with 1 mg/mL α-amylase for 1h at 37ºC then with 2 mg/mL Pronase for overnight at 60ºC to remove trapped high-molecular weight glycogen and peptidoglycan-associated proteins, respectively. Further PG preparation steps were performed as previously described (*42*). Briefly, muropeptides were released from PG by the muramidase cellosyl (Hoechst, Frankfurt am Main, Germany), reduced by sodium borohydride, and separated on a 250 × 4.6 mm 3 µm Prontosil 120-3-6C18 AQ reversed phase column (Bischoff, Leonberg, Germany). The eluted muropeptides were detected by their absorbance at 205 nm. The PG composition from exponentially and stationary growing cells was analyzed in 2 biological replicates. Muropeptides were assigned according to their retention times of the known muropeptides from *E. coli* and *A. baumannii* (*43*), and quantified using the Laura software (Lab Logic Systems).

*MS/MS analysis*

New muropeptide fractions with retention times other than standard muropeptides were collected and analyzed by MS/MS as previously described (*44*). LTQ-FT MS analyses revealed the presence of D-lysine in the collected muropeptide fraction c-e (Fig. S7). Table S1 shows the proposed structures, and the theoretical neutral and measured neutral atomic mass units. Muropeptide fractions a and b were also acetylated using an established protocol (*17*) prior MS/MS analysis, to confirm the number of amino groups.

*Construction of A. baumannii mutant and complement strains*

The primers used in this study are listed in Supplementary Table S5. Mutants were constructed as described previously(*45*). Briefly, an antibiotic resistance cassette was amplified with primers pair P1_kan Fwd-Rev (Integrated DNA Technologies) with homology to the flanking regions of the *racK* gene with additional 3′ 18–25 nucleotides of homology to the FRT site-flanked kanamycin resistance cassette from plasmid pKD4. This PCR product was electroporated into competent Ab17978 carrying pAT04, which expresses the RecAB recombinase (*45*). Mutants were selected on 10 μg per ml kanamycin, and integration of the resistance marker was confirmed by PCR. To remove the kanamycin resistance cassette, electrocompetent mutants were transformed with pAT03 plasmid, which expresses the FLP recombinase. To create Ab17978 RacK+ strain, the *racK* gene was cloned into pSH vector and under an arabinose inducible promotor, then the pSH_RacK plasmid was electroporated into Ab17978 ΔRacK strain. The *racK* gene was introduced into UPAB1 via a four-parent conjugal strategy as described by Kumar et al.(*46*). Briefly, 100 μl of stationary cultures normalized to an OD600 of 2.0 of each recipient strain, HB101(pRK2013), EC100D(pTNS2), and EC100D containing the pUC-miniTn7-racK plasmid were added to 600 µl of warm LB broth. Each suspension was washed twice by centrifugation at 7,000 × g followed by resuspension of the bacterial pellet in 1 ml of warm LB broth. On the final wash, the bacterial pellet was resuspended in 25 µl of LB broth, and the suspension was spotted on a prewarmed low salt LB agar plate and incubated overnight at 37°C. The bacteria were scraped from the plate and resuspended in 1 ml of LB broth, vortexed, and serial dilutions were plated on L agar plates supplemented with chloramphenicol to select against *E. coli* strains and kanamycin or zeocin to select for *A. baumannii* strains that had received the mini-Tn7 constructs. To verify that mini-Tn7 had successfully transposed downstream of the *glmS2* gene, it is amplified by PCR using primers pair UPAB1_check Fwd-Rev and the products were verified by sequencing. The strains were saved as Ab17978 RacK+ and UPAB1 RacK+.

*Protein expression and purification*

The *amaD* gene was PCR amplified from *Pseudomonas putida* and was cloned in pET22b+ vector void of the *pelB* sequence using Hi-Fi DNA Assembly mix (NEB). The pET22b+_amaD vector was electroporated into *E. coli* Rosetta II (Invitrogen) and selected on carbenicillin. The bacteria were grown in LB to an OD600 of 0.6 and AmaD protein expression is induced by addition of IPTG 0.5 mM for 3h at 30°C. Cells were harvested and lysed with two rounds of a cell disruptor using 35 k.p.s.i. (Constant System ltd., Kennesaw, GA). Cell lysates were clarified at 10,000 rpm for 10 min, then was passed over a nickel-nitrilotriacetic acid-agarose column Ni-NTA (Gold Bio, St. Louis, MO). Protein was eluted with 300 mM of Imidazole, buffer exchanged using Sephadex G25 PD10 column and stored in 50mM Tris-HCl pH 8.0, 150 mM NaCl at -80°C.

*P. aeruginosa* Tse1 protein was obtained as described (*18*). LdtA from *Vibrio cholerae* was obtained as described (*22*).

*AmaD enzymatic assay*

3 mL of bacteria cultures grown for 20 h were centrifuged at 10,000 x g and 0.5 mL of supernatants were collected. The supernatants were deproteinized using Amicon Ultra-0.5 mL Centrifugal Filters with 3,000 MWCO. Samples was incubated with 0.4 mg/ml of AmaD for 1 h at 37°C. Hydrogen peroxide co-product was then measured using Hydrogen Peroxide Assay Kitassay (Abcam) as instructed by manufacturer.

*Muropeptide isolation*

TetraTetra and TetraTri-D-Lys were purified by HPLC on an Aeris peptide column (250 × 4.6 mm; 3.6 µm particle size; Phenomenex, USA), concentrated and desalted using a water-methanol gradient on the same column prior to MS analysis. TetraTetra was obtained from muramidase treatment of *Escherichia coli* sacculi (*47*). Chromatographic analyses of muropeptides were performed on AQUITY Ultra Performance Liquid Chromatography (UPLC) BEH C18 column (130 Å, 1.7 µm, 2.1 mm by 150 mm; Waters), and peptides were detected at Abs. 204 nm using ACQUITY UPLC UV-Visible Detector. Muropeptides were separated using a linear gradient from buffer A (0.1% of Formic acid in water) to buffer B (0.1% of Formic acid in acetonitrile) in 18 min, and flow 0.25 ml/min. Muropeptide identity was confirmed by MS/MS analysis, using a Xevo G2-XS QTof system (Waters Corporation, USA). Quantification of muropeptides was based on their relative abundances (relative area of the corresponding peak).

*In vitro reaction*

TetraTri-D-Lys was obtained from mixing 10 µg of TetraTetra with 20 mM of D-Lys and 1µM of LdtA for 4h in Tris pH8 20 mM. Peptidoglycan digestion was analyzed over time (0.5, 1 and 5 min). 0.01 mg/ml of purified Tse1 enzyme from *P. aeruginosa* at 37°C in buffer 20 mM Tris HCl pH 8.0 was used in the different reactions. The activities for each substrate concentration and time were assessed in triplicate. Individual muropeptides were quantified from their integrated areas using samples of known concentration as standards. Reactions were terminated by inactivation at 100°C for 10 min and centrifuged at 14,000 x g for 30 min to discard the coagulated

Tse1. Next, Tse1-treated muropeptides were analyzed by UPLC. Determination of the extent of Tse1-dependent degradation was performed by comparing the integration areas with respect to a non-treated sample.

*T6SS competition assay*

Competition assays were performed as previously described (*48*). Briefly, predator and prey overnight cultures were pelleted, washed three times in fresh LB, and resuspended at an OD600nm of 1.0. The cultures were mixed at a predator:prey ratio of 5:1 (PA01:Ab or M2:Ab) or 10:1 (Serratia:Ab), and 10 µl drops were spotted on a LB 3% agar plate. After 4 h at 37°C, the spots were harvested, resuspended in 0.7 mL of LB broth, serially diluted and plated on rifampicin or gentamycin LB agar plates in order to determine the number of surviving prey cells. In parallel, the number of surviving PA01 predator was also enumerated on irgasan LB agar plates.

*Murine model of A. baumannii acute pneumonia*

All infection experiments were approved by the Vanderbilt University Institutional Animal Care and Use Committee and are in compliance with guidelines set by the Animal Welfare Act, the National Institutes of Health, and the American Veterinary Medical Association. Lung infection experiments were performed essentially as previously described(*32*). Briefly, prior to infection, overnight cultures were subcultured 1:100 in 30 mL of liquid LB media and incubated at 37 °C with shaking. Bacteria in mid-exponential phase growth were harvested by centrifugation, washed twice with PBS, and resuspended in PBS to the same concentration. Anesthetized 7-9 week-old C57BL/6 mice (Jackson Laboratories) were inoculated intranasally with 4-6 x $10^8$ CFU of Ab17978, UPAB1, or their respective isogenic mutants in 35 µL PBS. Lungs, livers, spleens, kidneys, hearts and blood were aseptically harvested from mice euthanized at 36 h post-infection. Organs were homogenized and then serially diluted and plated to LB agar to determine bacterial burdens.

*DAO antimicrobial in vitro assay*

Three ml of bacteria cultures grown for 20 h were centrifuged at 10,000 x g and 900 µL of supernatants were collected. One hundred microliters of fresh LB was added as well as purified porcine kidney DAO and Catalase (Millipore Sigma) to 50 µg/ml and 25 µg/ml respectively. The bacterial pellets were resuspended and added to the supernatant-LB-DAO or DAO/Catalase mixtures to an OD600 of 0.005. After 4.5 h of incubation at 37°C, bacteria were serial diluted and spotted on LB agar plates for enumeration.

*Genome Data*

Genomic data used in this study were obtained from the NCBI RefSeq data base (version 87, accessed March 13, 2018). All 3052 Acinetobacter spp. genome assemblies together with the corresponding coding and protein sequences were downloaded into a local database and integrated in this analysis. To reduce the computational burden for tree reconstruction, we compiled a core set of 232 Acinetobacter spp. encompassing all described international clone types of *A. baumannii* as well as type and reference strains of 53 validly named species.

*Phylogenetic Profiling and Synteny Analysis*

For all protein sequences, orthologous groups (OGs) across the 3052 genomes were inferred using a tandem of OMA standalone v. 2.2.0 [10.1093/nar/gkx1019] and HaMStR v. 13.2.9 [https://github.com/BIONF/hamstr, 10.1186/1471-2148-9-157] as previously described [10.1080/21505594.2018.1558693]. The synteny of *racK* and its two flanking genes was inferred with the genomic loci of the corresponding orthologs using the gene order and orientation of the strain ATCC 17978 as a reference.

*Phylogenetic Tree reconstruction*

The protein sequences of *racK* and its flanking genes in the core set were aligned for each orthologous group individually with MAFFT-linsi v7.407 [10.1093/nar/gkt389]. The resulting multiple sequence alignment together with the corresponding coding sequences served then as input for PAL2NAL v.14 [10.1093/nar/gkl315] (with option '-codontable 11') to generate corresponding codon alignments. Individual or concatenated MSAs were then used for a maximum likelihood tree reconstruction with IQ-Tree v1.6.12 [10.1093/nar/gkw256] performing 1000 non-parametric bootstrap replicates [10.1093/molbev/msx281]. The best-fitting substitution model was determined with the model-testing routines implemented into IQ-Tree [10.1038/nmeth.4285]. Topology tests were performed using the weighted and unweighted Shimodaira-Hasegawa test [10.1093/oxfordjournals.molbev.a026201] as implemented in IQ-Tree.

*MLST classification and IC assignments*

We predicted for each of the 3052 Acinetobacter spp. strains the sequence type with two different MLST schemes, Oxford [10.1128/JCM.43.9.4382-4390.2005] and Pasteur [10.1371/journal.pone.0010034], that we obtained from the pubmlst website (http://pubmlst.org/abaumannii/) using MLSTcheck v2.1.17 [10.21105/joss.00118]. Strains of A. baumannii were assigned to an international clone type (IC) if predicted sequence types and IC were unambiguously linked in the literature [10.1016/j.ijantimicag.2019.03.019, 10.1016/j.meegid.2019.103986, 10.1016/j.watres.2018.04.057, 10.1038/s41426-018-0127-9, 10.1111/1462-2920.13931, 10.1186/1471-2180-13-234, 10.1371/journal.pone.0010034, 10.1371/journal.pone.0153014, 10.1371/journal.pone.0179228, 10.3389/fmicb.2019.00930, 10.1186/s13059-015-0701-6]

*Statistical analysis*

All statistical analyses were performed using GraphPad Prism (GraphPad Software Inc., La Jolla, CA). For the murine acute pneumonia model, data were log transformed and analyzed for Gaussian distribution using the D'Angostino-Pearson omnibus normality test. For normally distributed datasets, Student's unpaired t tests were used. For datasets with unknown distribution type, nonparametric Mann-Whitney U tests were used.

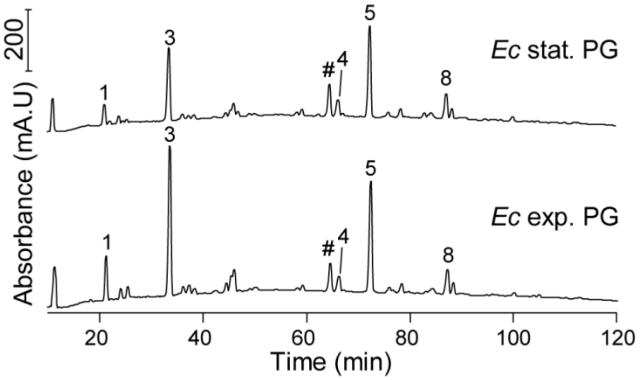


**Fig. S1. Stationary *E. coli* cells do not modify their PG.** Muropeptide profiles of exponential (exp.) and stationary (stat.) phase *E. coli* BW25113 (*Ec*) cells. Main muropeptides are labelled and their structures are shown in Fig. S2.

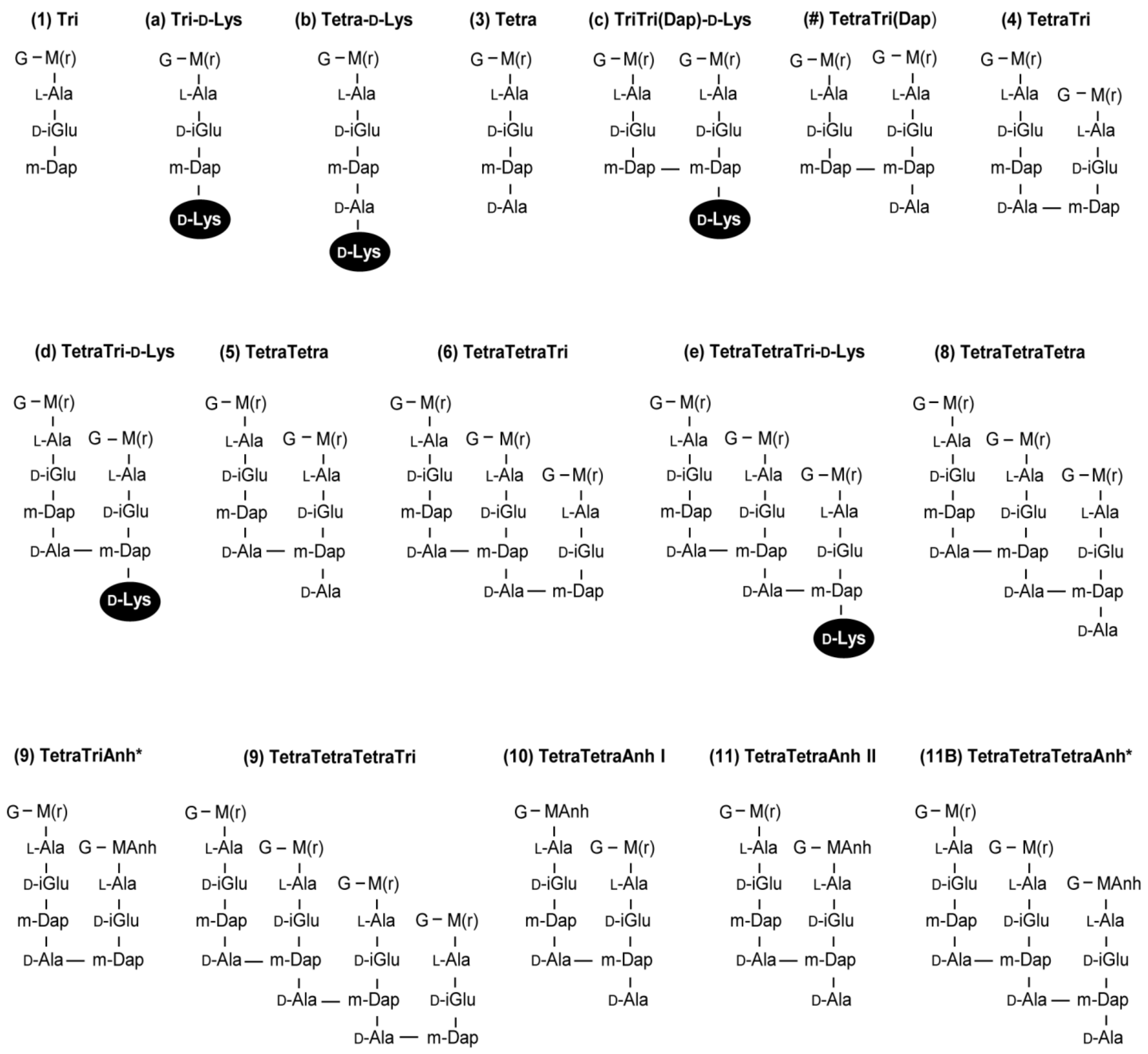


**Fig. S2. Proposed muropeptide structures of the chromatograms shown in Figs. 1, 2C, 3B, S1 and S4.** M(r), *N*-acetylmuramitol; G, *N*-acetylglucosamine. MS analysis of peak 9 reavealed a composition of two muropeptides (~70% d43Anh and ~30% p4443). * position of the anhydro-group is unknown. Peak 0 is generated by acid hydrolysis of unused lipid II or glycan chains ends carrying the C55-PP moiety. m = monomer, d = dimer, t = trimer, p = polymer. 3,4,5 = number of amino acids in the peptide stem. Anh = anhydro-group.

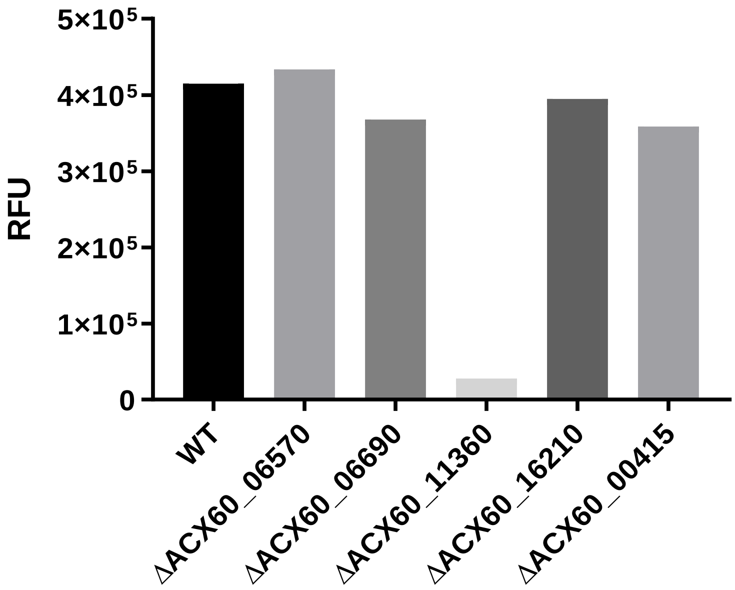


**Fig. S3. Identification of racemase responsible for D-Lysine production in Ab17978.** AmaD assay described in the Methods section was used to quantify D-Lys concentrations of stationary phase culture supernatants of *A. baumannii* 17978 strains individually lacking possible racemases.

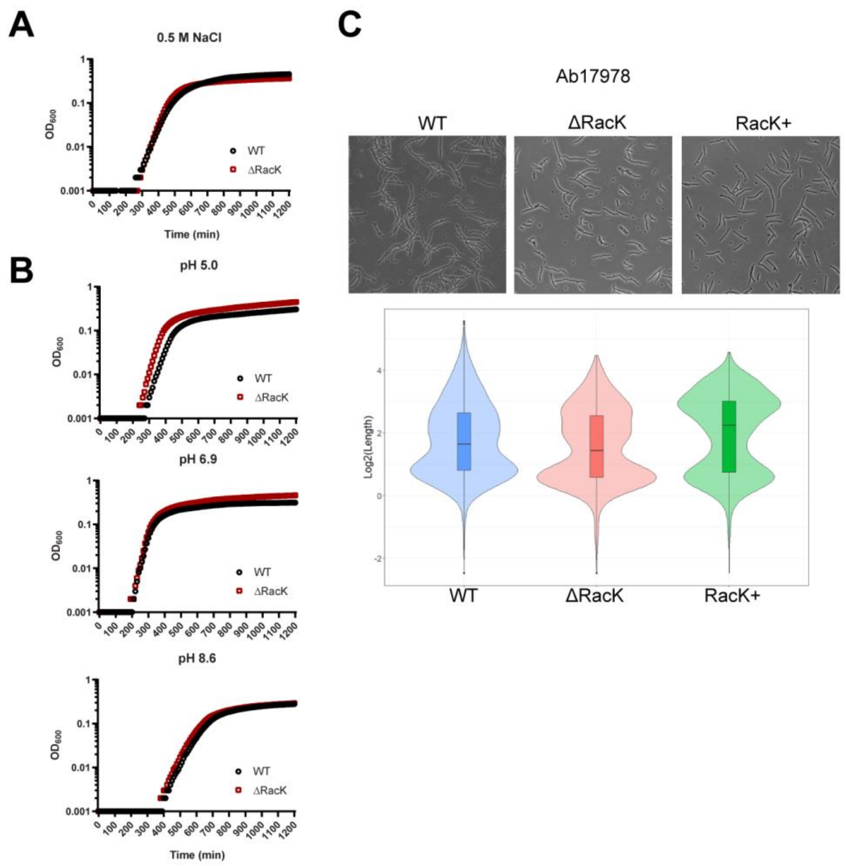


**Fig. S4. PG modification in Ab17978 does not have notable impact on its response to osmotic (A) and pH stress (B) or bacteria morphology (C). (A)** Growth curve of Ab17978 WT or ΔRacK grown in LB supplemented with 0.5M NaCl. **(B)** Growth curve of Ab17978 WT or ΔRacK growth in LB buffered at pH 5.0, 6.9 or 8.6. **(C)** Ab17978 WT, ΔRacK or RacK+ bacteria cells were observed under phase contrast microscopy. Cell length of each strain was obtained by using cell segmentation SuperSegger algorithm (Stylianidou et al. 2016) on 10 microscopy images. Violin and box plots represent length distribution of 800+ bacteria cells in log2 scale.

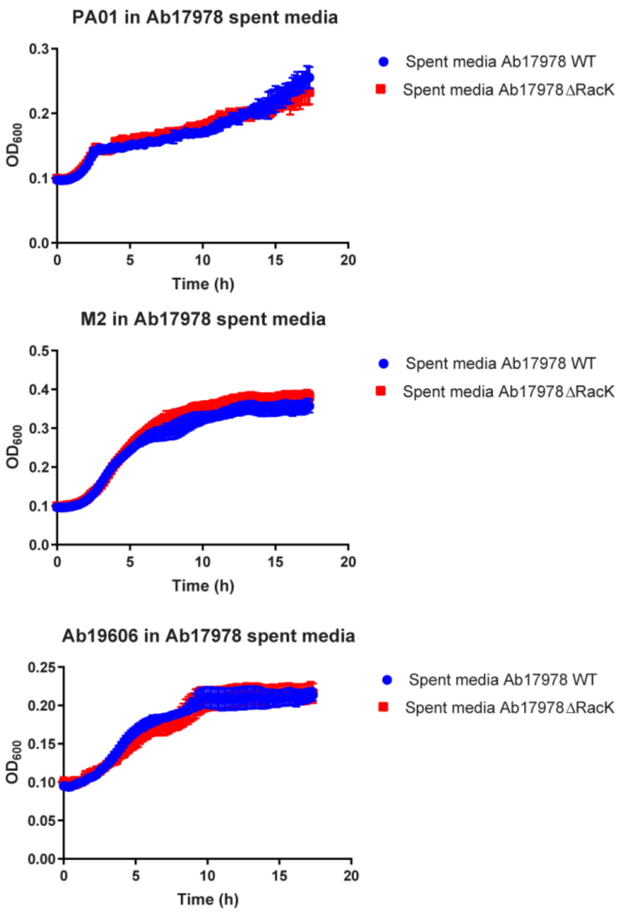


**Fig. S5. Concentration of secreted D-Lys by Ab17978 is not enough to induce inter-bacterial growth inhibition in liquid culture.** Stationary culture supernatants of Ab17978 WT and ΔRacK were use as growth media of *P. aeruginosa* PA01, *A. nosocomialis* M2 and *A. baumannii* Ab19606. Data are representative of 3 biological replicates.

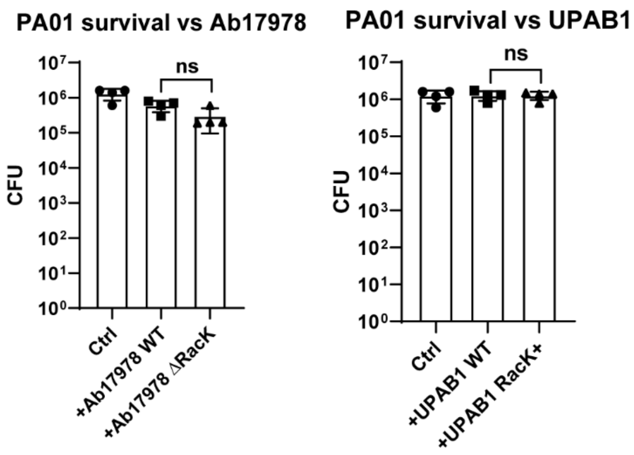


**Fig. S6.** ***P. aeruginosa*** **PA01 (PA01) growth is unaffected by co-incubation with Ab17978 or UPAB1 during T6SS killing assays shown in Fig. 2.** Bar graphs represent the mean ± SD of 4 biological replicates. Statistical analyses were performed using the unpaired Student's t test, ns: non significative.

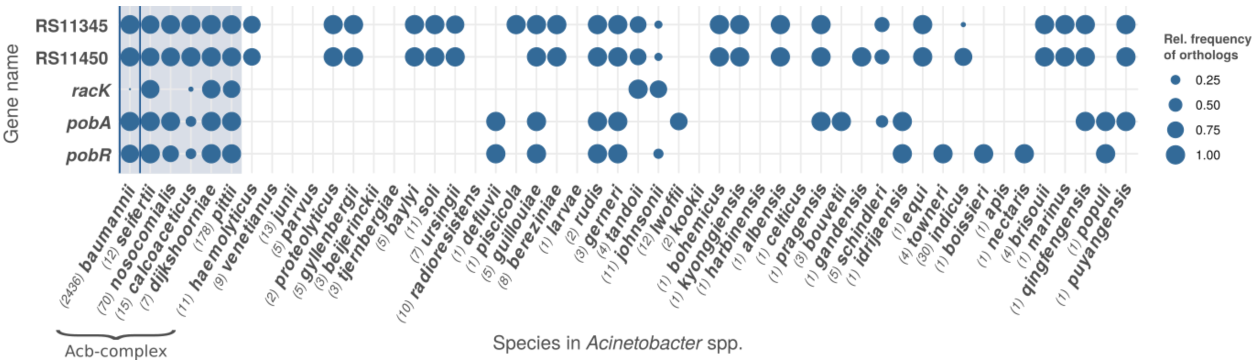


**Fig. S7A. Presence/absence pattern of *racK* and its flanking genes in the genus *Acinetobacter*.** The phylogenetic profile reveals the occurrence of orthologs to *racK* and of its two flanking genes up- and downstream, respectively, across 3,052 *Acinetobacter spp.* genomes. The information was summarized on the species level with the number of genomes per species given in parentheses. A blue dot indicates the presence of an ortholog in the respective species. The dot diameter is proportional to the relative frequency of genomes that harbour an ortholog. In total, only 7 species of *Acinetobacter* harbour an ortholog including *A. tandoii* and *A. johnsonii*. Within the Acb-complex *racK* orthologs were found in only 7% of all *A. baumannii* strains (cf Figure 7), 10% of *A. calcoaceticus* and none of *A. nosocomialis*. In contrast, it is almost ubiquitously present in *A. seiffertii*, *A. pittii*, and *A. dijkshoorniae*. The presence of orthologs to all five genes is restricted to the clade constituting the Acb-complex. The profile was visualized using PhyloProfile v. 1.1.2 [10.1093/bioinformatics/bty225].

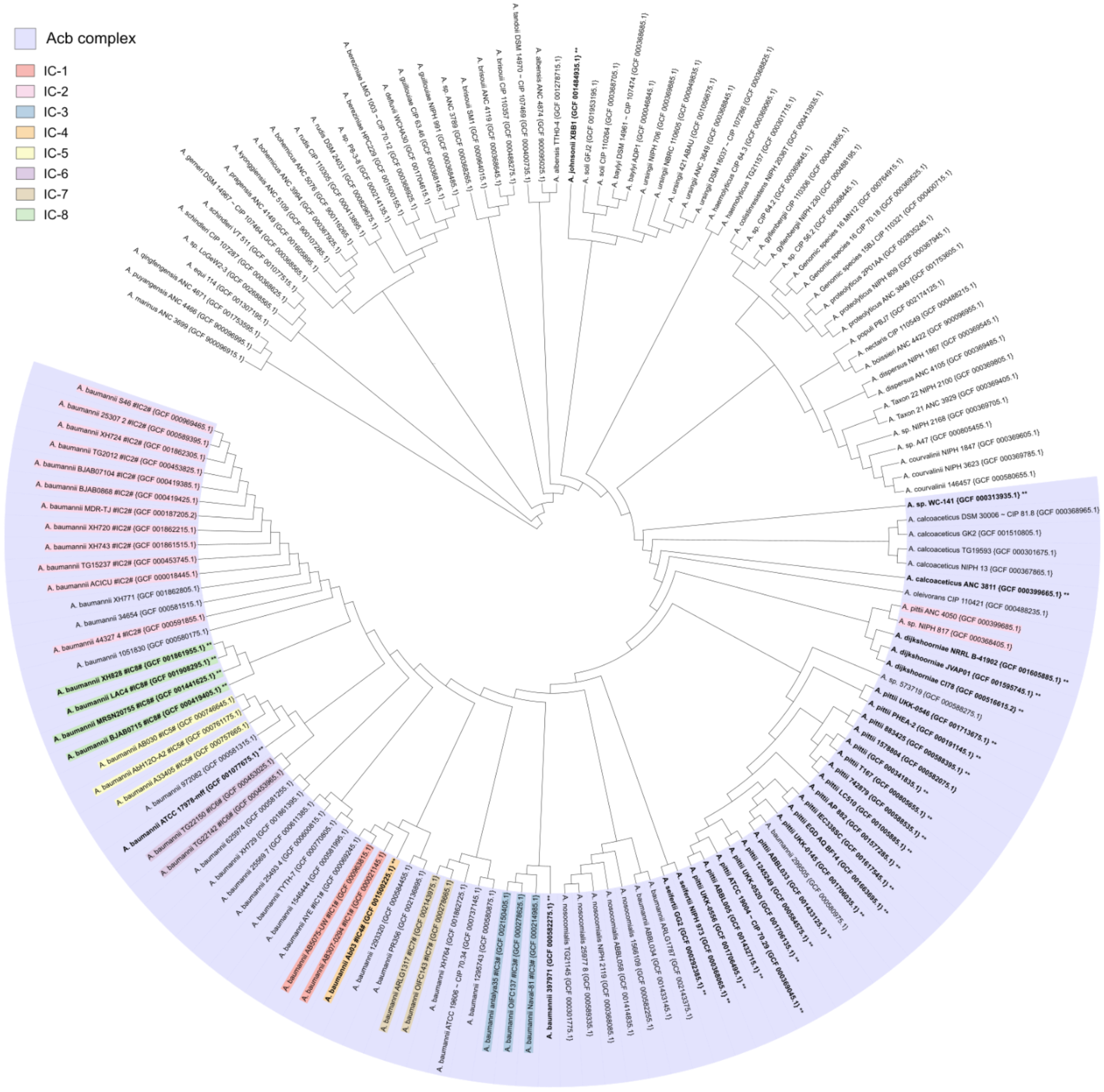


**Fig. S7B. Maximum-Likelihood phylogeny of racK's genomic neighborhood across Acinetobacter spp.** The tree was reconstructed under the best fitting model TIM+F+I+G4 and was inferred from concatenated codon alignments of racK's 4 flanking genes (ACX60_RS11345, ACX60_RS11350, ACX60_RS11360, and ACX60_RS11365) that we traced across 232 genomes. Leaf labels comprise species names, strain names and identifiers, as well as RefSeq assembly accessions (in curly brackets). Clone Type (IC) affiliation of an A. baumannii strain is indicated by label background colors (cf. figure legend). For visualization, the branch lengths have universally been set to 1. Branch labels specify bootstrap supports. Strains harboring a racK ortholog (bold faced tip labels) are distributed within the phylogeny of the Acinetobacter calcoaceticus/baumannii (Acb-) complex (blue background).

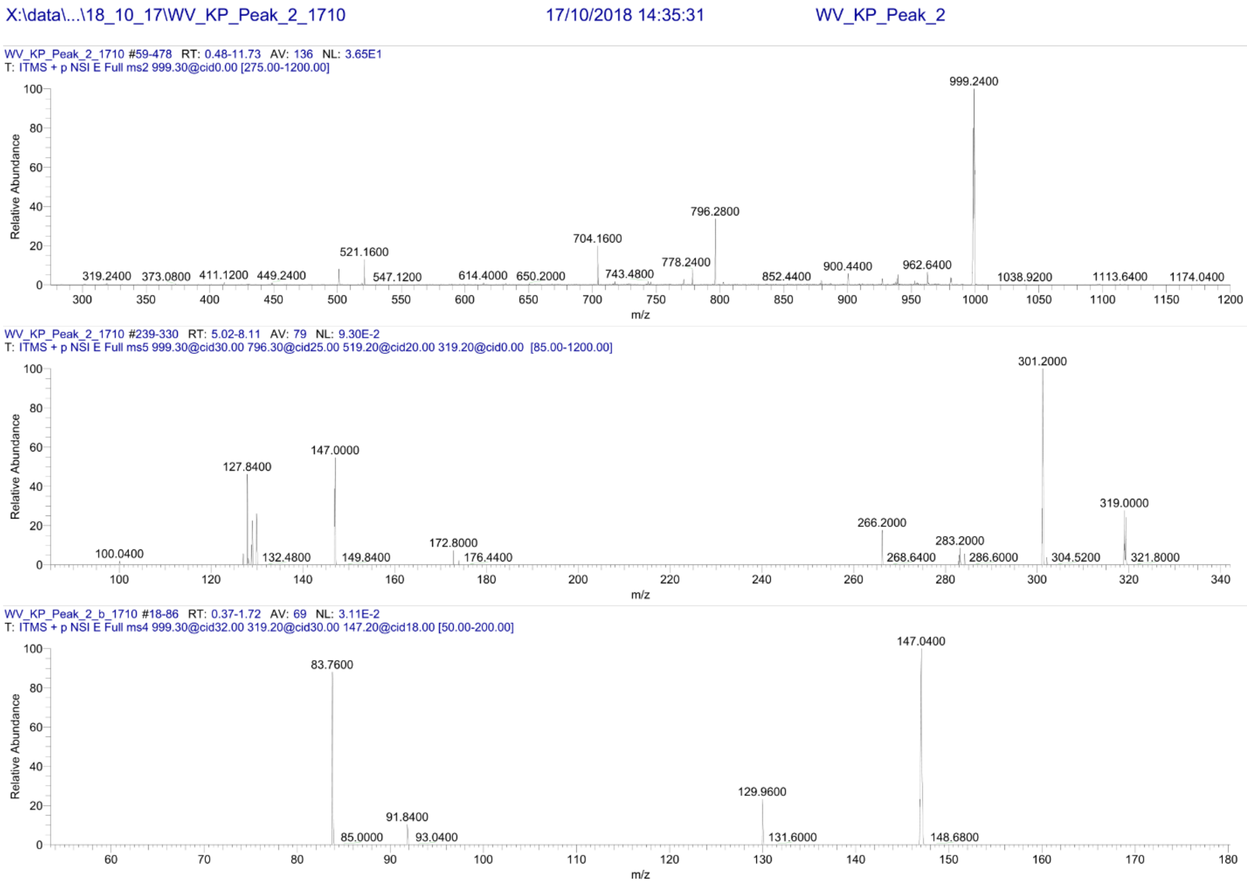


**Fig. S8A.** Mass spectrometry analysis of peak a (Tri-D-Lys) as presented in Fig. 1

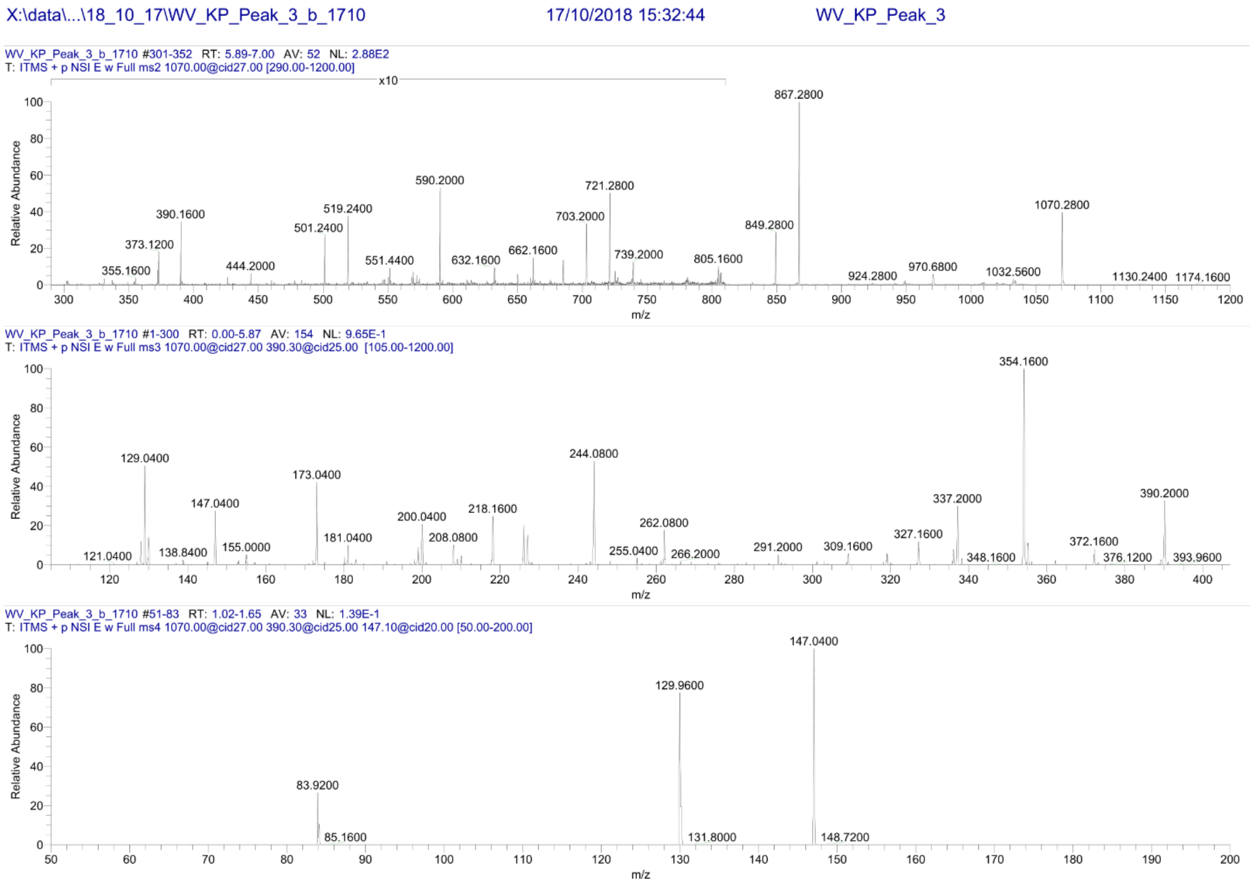


**Fig. S8B.** Mass spectrometry analysis of peak b (Tetra-D-Lys) as presented in Fig. 1

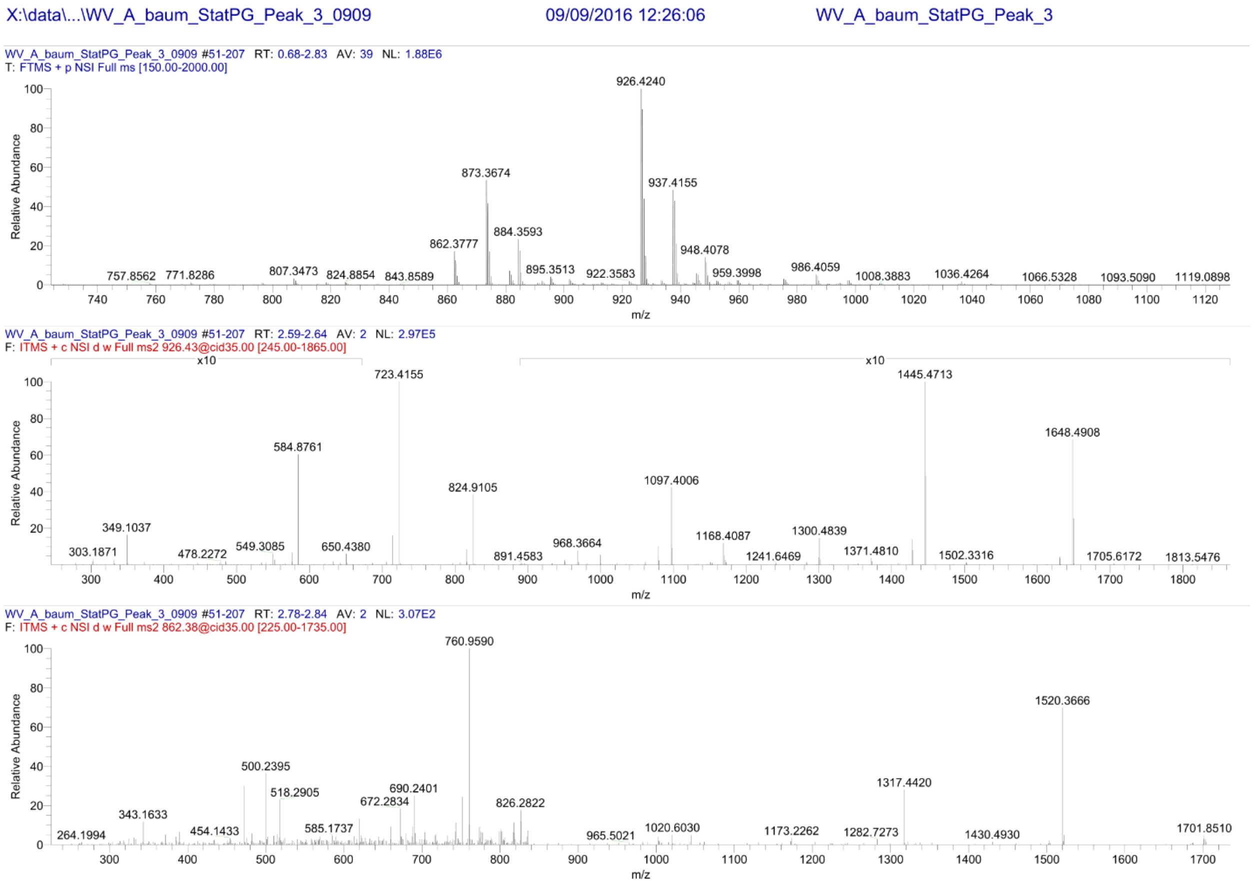


**Fig. S8C.** Mass spectrometry analysis of peak c (TriTri(Dap)-D-Lys) as presented in Fig. 1

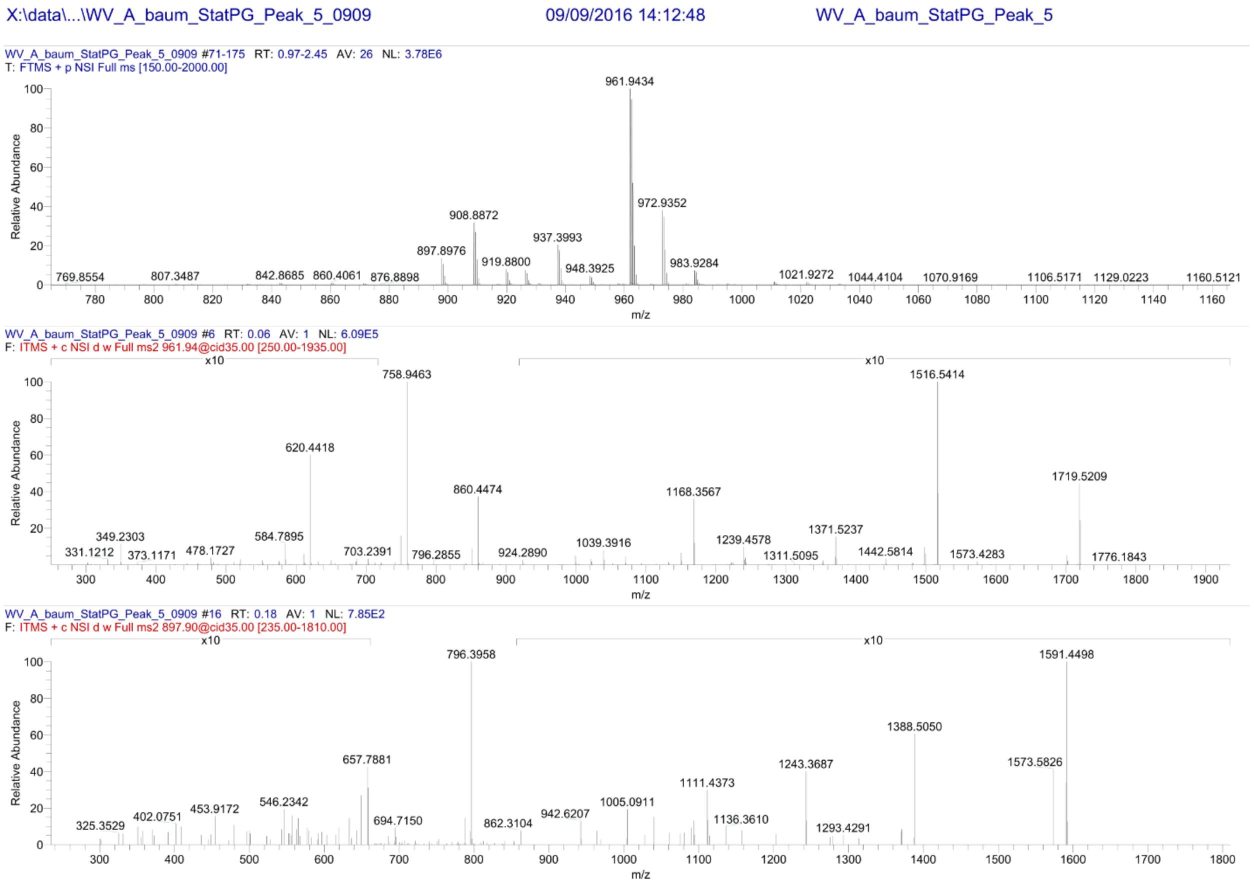


**Fig. S8D.** Mass spectrometry analysis of peak d (TetraTri-D-Lys) as presented in Fig. 1

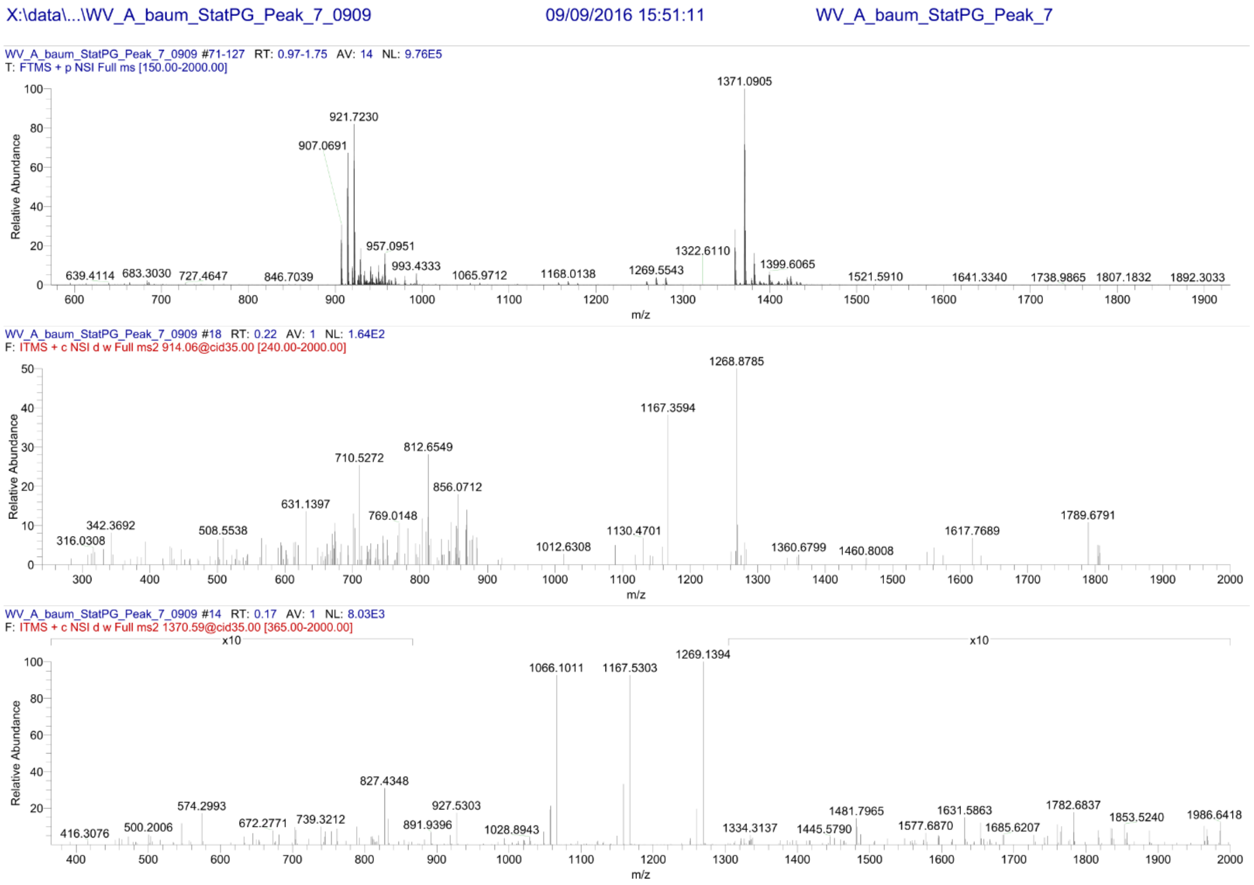


**Fig. S8E.** Mass spectrometry analysis of peak e (TetraTetraTri-D-Lys) as presented in Fig. 1

**Table S1.** Reduced D-Lys containing muropeptides from stationary *A. baumannii* 17978 cells in HPLC fractions detected by LTQ- FT MS.

| Label | Proposed structure(s)[1] | Retention time (min) | Theoretical neutral mass (Da) | Measured neutral mass (Da) |
|---|---|---|---|---|
| **1** | **Tri** | 21.2 | | |
| **a** | **Tri-D-Lys** | 24.4 | 998.4656 | 998.3662 |
| **b** | **Tetra-D-Lys** | 32.5 | 1069.5027 | 1069.4276 |
| **3** | **Tetra** | 33.3 | | |
| **c** | **TriTri(Dap)-D-Lys** | 59.0 | 1850.8256 | 1850.8324 |
| **4** | **TetraTri** | 64.1 | | |
| **d** | **TetraTri-D-Lys** | 66.4 | 1921.8627 | 1921.8712 |
| **5** | **TetraTetra** | 72.0 | | |
| **6** | **TetraTetraTri** | 82.4 | | |
| **e** | **TetraTetraTri-D-Lys** | 84.2 | 2845.2598 | 2845.7044 |
| **8** | **TetraTetraTetra** | 88.1 | | |
| **9** | **TetraTetraTetraTri/ TetraTriAnh[2]** | 96.4 | | |
| **10** | **TetraTetraAnh I** | 100.6 | | |
| **11** | **TetraTetraAnh II** | 102.1 | | |
| **11B** | **TetraTetraTetraAnh** | 112.2 | | |

[1] Nomenclature of muropeptides according to Glauner (1988). Muropeptides were assigned according to their retention times, which were identical to known, unmodified muropeptides from *E. coli*. Muropeptides were numbered according to Boll *et al*., 2016. D-Lys containing muropeptides (a-e) were confirmed by MS/MS-analysis.
[2] MS analysis of peak 9 reavealed a mixture of two muropeptides (~70% TetraTriAnh and ~30% TetraTetraTetraTri).

**Table S2.** Exponential and Stationary peptidoglycan composition of *A. baumannii* strains (Quantification of the chromatograms shown in Fig. 1A)

| | | Relative peak area (%)[2] | |
|---|---|---|---|
| **Label** | **Muropeptide[1]** | **17978 expo.** | **17978 stat.** |
| **1** | **Tri** | 3.0 | 2.1 |
| **a** | **Tri-D-Lys** | 2.1 | 10.5 |
| **b** | **Tetra-D-Lys** | 0.0 | 10.0 |
| **3** | **Tetra** | 19.6 | 10.7 |
| **c** | **TriTri(Dap)-D-Lys** | 1.0 | 3.3 |
| **4** | **TetraTri** | 6.1 | 4.3 |
| **d** | **TetraTri-D-Lys** | 0.9 | 8.1 |
| **5** | **TetraTetra** | 37.5 | 28.2 |
| **6** | **TetraTetraTri** | 1.6 | 1.5 |
| **e** | **TetraTetraTri-D-Lys** | 2.6 | 2.5 |
| **8** | **TetraTetraTetra** | 17.1 | 12.9 |
| **9** | **TetraTriAnh (~70%)[3]** | 3.2 | 2.3 |
| **9** | **TetraTetraTetraTri (~30%)[3]** | 1.4 | 1.0 |
| **10** | **TetraTetraAnh I** | 0.5 | 0.2 |
| **11** | **TetraTetraAnh II** | 0.7 | 0.5 |
| **11B** | **TetraTetraTetraAnh** | 2.8 | 1.9 |
| **Monomers** | | 24.7 | 33.3 |
| **Dimers** | | 49.8 | 47.0 |
| **Trimers** | | 24.1 | 18.7 |
| **Tetramers** | | 1.4 | 1.0 |
| **% peptides in crosslinkage** | | 75.3 | 66.7 |
| **Average chain length[4]** | | 31.9 | 46.1 |
| **D-Lys containing muropeptides** | | 6.5 | 34.4 |

[1] Muropeptides 1-11B are numbered according to Boll *et al*., 2016 and D-Lys containing muropeptides are labelled a-e. [2] The relative peak areas were estimated as percentage of all known peaks. [3] MS analysis of peak 9 revealed a composition of two muropeptides (~70% TetraTriAnh and ~30% TetraTetraTetraTri). [4] The chain length is given in disaccharide unit calculated from the % of anhydro groups in the compound.

**Table S3.** Exponential and Stationary peptidoglycan composition of *E. coli* (Quantification of the chromatograms shown in Fig. S1)

| | | Relative peak area (%)[2] | |
|---|---|---|---|
| **Label** | **Muropeptide[1]** | **E. coli expo.** | **E. coli stat.** |
| **1** | **Tri** | 9.15 | 6.72 |
| **3** | **Tetra** | 39.09 | 29.34 |
| ***** | **TetraTri(Dap)** | 7.80 | 12.01 |
| **4** | **TetraTri** | 4.86 | 7.43 |
| **d** | **TetraTri-D-Lys** | 0.9 | 8.1 |
| **5** | **TetraTetra** | 31.17 | 34.15 |
| **6** | **TetraTetraTri** | 0.00 | 0.00 |
| **8** | **TetraTetraTetra** | 7.92 | 10.35 |
| **Monomers** | | 48.24 | 36.06 |
| **Dimers** | | 43.83 | 53.59 |
| **Trimers** | | 7.92 | 10.35 |
| **% peptides in crosslinkage** | | 51.76 | 63.94 |

**Table S4.** Stationary peptidoglycan composition of *A. baumannii* strains (Quantification of the chromatograms shown in Fig. 2C)

| | | Relative peak area (%)[2] | | |
|---|---|---|---|---|
| **Label** | **Muropeptide[1]** | **17978** | **17978 ΔRacK** | **17978 RacK+** |
| **1** | **Tri** | 2.2 | 4.1 | 3.9 |
| **a** | **Tri-D-Lys** | 11.5 | 0.6 | 9.9 |
| **b** | **Tetra-D-Lys** | 10.4 | 0.3 | 5.4 |
| **3** | **Tetra** | 10.6 | 12.5 | 12.6 |
| **c** | **TriTri(Dap)-D-Lys** | 3.5 | 0.8 | 2.3 |
| **4** | **TetraTri** | 4.7 | 4.0 | 6.0 |
| **d** | **TetraTri-D-Lys** | 9.2 | 2.6 | 7.5 |
| **5** | **TetraTetra** | 21.3 | 30.7 | 26.4 |
| **6** | **TetraTetraTri** | 1.4 | 2.1 | 1.9 |
| **e** | **TetraTetraTri-D-Lys** | 4.8 | 2.1 | 3.5 |
| **8** | **TetraTetraTetra** | 11.1 | 23.0 | 12.1 |
| **9** | **TetraTriAnh (~70%)[3]** | 3.1 | 6.7 | 2.0 |
| **9** | **TetraTetraTetraTri (~30%)[3]** | 1.3 | 2.9 | 0.9 |
| **10** | **TetraTetraAnh I** | 0.8 | 3.0 | 1.6 |
| **11** | **TetraTetraAnh II** | 1.2 | 1.2 | 1.4 |
| **11B** | **TetraTetraTetraAnh** | 2.9 | 3.2 | 2.7 |
| **Monomers** | | 34.7 | 17.6 | 31.8 |
| **Dimers** | | 43.8 | 49.1 | 47.2 |
| **Trimers** | | 20.2 | 30.5 | 20.1 |
| **Tetramers** | | 1.3 | 2.9 | 0.9 |
| **% peptides in crosslinkage** | | 65.3 | 82.4 | 68.2 |
| **Average chain length[4]** | | 28.5 | 15.4 | 29.5 |
| **D-Lys containing muropeptides** | | 39.4 | 6.5 | 28.5 |

[1] Muropeptides 1-11B are numbered according to Boll *et al*., 2016 and D-Lys containing muropeptides are labelled a-e. [2] The relative peak areas were estimated as percentage of all known peaks. [3] MS analysis of peak 9 revealed a composition of two muropeptides (~70% TetraTriAnh and ~30% TetraTetraTetraTri). [4] The chain length is given in disaccharide unit calculated from the % of anhydro groups in the compound.

**Table S5.** Bacterial strains and plasmids used in this study

| **Bacterial strains** | **Description** |
|---|---|
| Ab17978 | *Acinetobacter baumannii* strain ATCC17978 devoid of pAB3 plasmid. Spontaneous Rifampin resistant. (Weber et al.) |
| Ab17978 ΔRacK | Ab17978 *racK* (*acx60_11360*) deletion mutant. Spontaneous Rifampin resistant |
| Ab17978 ΔACX60_06570 | Ab17978 *acx60_06570* deletion mutant |
| Ab17978 ΔACX60_06690 | Ab17978 *acx60_06690* deletion mutant |
| Ab17978 ΔACX60_16210 | Ab17978 *acx60_11210* deletion mutant |
| Ab17978 ΔACX60_00415 | Ab17978 *acx60_00415* deletion mutant |
| Ab17978 RacK+ | Ab17978 ΔRacK with pSH_RacK plasmid |
| UPAB1 | *Acinetobacter baumannii* strain UPAB1 (Di venanzio et al., 2019) |
| UPAB1 RacK+ | *Acinetobacter baumannii* strain UPAB1 with chromosomal insertion of racK and its 300 bp upstream region |
| Ab19606 | *Acinetobacter baumannii* strain ATCC19606 |
| M2 | *Acinetobacter nocosomialis* strain M2 with pVRL2 plasmid. Gentamycin resistant |
| PA01 | *Pseudomonas aeruginosa* strain PA01 |
| BW25113 | *Escherichia coli* strain BW25113 |
| Serratia | *Serratia marcescens* strain 66262 |

| **Plasmids** | **Description** |
|---|---|
| pET22b_AmaD | *amaD* gene from *Pseudomonas putida* cloned into pET22b+ plasmid |
| pBAD_RacK | *racK* gene from Ab17978 cloned into an in-house modifier version of the arabinose inducible vector pBAD |
| pUC-miniTn7-racK | *racK* with its 300 bp upstream region from Ab17978 cloned into a pUC18 vector |
| pVRL2 | As described in Lucidi et al. 2018, Gentamycin resistant |

**Table S6.** Primers used in this study

| Primers | Sequence |
|---|---|
| AB RacK Fwd | ATAAAACAAAGTTTCGGATG |
| AB RacK Rev | TCCAGCCTACACAATCGCGAGTTTTTTAATCTTTCCTGG |
| CD RacK Fwd | TAAGGAGGATATTCATATGTAAACTTTAGGTGAATTGATAAG |
| CD RacK Rev | ATGCTGCGCATATTGTTCC |
| AB 06570 Fwd | ATTAGCTTAAGTGAGGCTTCG |
| AB 06570 Rev | TCCAGCCTACACAATCGCTATTTAAACCTTTTACTGGAC |
| CD 06570 Fwd | TAAGGAGGATATTCATATGTAAATACTCATCAACTTTTAATTTATTCTG |
| CD 06570 Rev | TTAGTTTAGGTCGTGTTGTCG |
| AB 06690 Fwd | TACTGAAGTTCGTGCCCGAGC |
| AB 06690 Rev | TCCAGCCTACACAATCGCCCTCATTTACTCCTTTGGGC |
| CD 06690 Fwd | TAAGGAGGATATTCATATGTAAAGTTTACTAAATAAAAAATCC |
| CD 06690 Rev | ATATCAGTTCAGATACCCAAG |
| AB 16210 Fwd | ATGGCATCCAAAGCATTGCGG |
| AB 16210 Rev | TCCAGCCTACACAATCGCGGCAGTACTAGAGAGTGTCGG |
| CD 16210 Fwd | TAAGGAGGATATTCATATGTAAGCTCTGTTGTAGGCATTCAG |
| CD 16210 Rev | AAGTTCTAAATCACGTGGGC |
| AB 00415 Fwd | ACAAACAACTTACGCCTAGAG |
| AB 00415 Rev | TCCAGCCTACACAATCGCGCTTTATCCTGTATTTCATAT |
| CD 00415 Fwd | TAAGGAGGATATTCATATGTAAAGCCAAGAAAAACGCCTAGCC |
| CD 00415 Rev | AAAATCTGGTCTAATGGCAGC |
| P1_kanR | AGCGATTGTGTAGGCTGGAGCTG |
| P2_kanR | CATATGAATATCCTCCTTAGTTCCTATTCCG |
| amaD Fwd | TAAGAAGGAGATATACATATGCATTGCCAGACCCTTGTC |
| amaD Rev | GTGGTGGTGGTGCTCGAGGTCGAAACGGGTCGCGCTGTA |
| pSH_racK Fwd | TAGCGGCCGCTGCAGGCCTATGAAACTTAAAACAAATTTTTC |
| pSH_racK Rev | TTAAGCGGCGGCATCGATCGTCAGTGGTGGTGGTGGTGGTGC |
| Prom_racK Fwd | ATGAGCTCACTAGTGGATCCAAAGTTTCGGATGTTTACCTC |
| Tn7_racK Rev | GAGGTACCGGGCCCAAGCTTTTACTCAGCGGTTGCTGCAAC |
| UPAB1_check Fwd | TAGAGCTATAAAAAGCCC |
| UPAB1_check Rev | TTGGCGAAGTCAGTAACTG |
